# Radiation therapy synergizes with mRNA vaccination to overcome microglia-mediated suppression of dendritic cell migration and T cell priming in glioblastoma

**DOI:** 10.64898/2026.09.18.752741

**Authors:** Amy J. Wisdom, Zachary J. Rogers, Fiona Chatterjee, Yufei Cui, Lizmarie Garcia-Rivera, Heidi Temple, Nouran S. Abdelfattah, Tobias Coombs, Richard Van, Cassandra Tafuri Del Vecchio, Hin Ka So, Taylor A. Heim, Grace Wolczanski, Christopher W. Mount, Vincent L. Butty, Stuart S. Levine, Kathleen Cormier, J. Christopher Love, Susan M. Kaech, Tyler Jacks, Forest M. White, Stefani Spranger

## Abstract

Glioblastomas (GBMs) are uniformly fatal brain tumors resistant to immunotherapy, yet the mechanisms driving immune dysfunction remain poorly understood. Here we identify a tissue-specific mechanism of dendritic cell (DC) dysfunction contributing to GBM immune evasion. We demonstrate that while systemic immunity depends on type 1 conventional DCs (cDC1s), GBMs preferentially accumulate cDC2s with impaired antigen-presenting capacity. Further, microglia-derived GAS6 and PROS1 signaling through the AXL receptor on DCs suppresses DC activation and, critically, migration to tumor-draining lymph nodes. Pharmacologic AXL inhibition enhances DC migration and consequently anti-tumor immunity. Importantly, radiation therapy activates cDC1-mediated endogenous immune responses, which synergizes with mRNA vaccination to promote complete and durable immune-mediated tumor regression. These findings establish DC-mediated immunity as a therapeutic target in brain tumors and reveal a synergistic mechanism for combining radiation with personalized cancer vaccines.

**Highlights:**

● The GBM TME is dominated by cDC2 responses, with poor accumulation of cDC1s and blunted systemic antitumor immunity
● Microglia suppress cDC migration and activation in the CNS via AXL signaling, and AXL blockade restores DC activation and migration
● Radiation therapy reactivates cDC1-mediated endogenous immunity against GBM
● Radiation therapy and tumor antigen-targeting mRNA vaccination achieve durable tumor cures in orthotopic and autochthonous GBM models

## Introduction

Glioblastomas (GBMs) are the most common primary brain tumors and remain lethal. Standard of care treatment for patients with GBM, which includes trimodality therapy with surgery, chemotherapy, and radiation therapy (RT)^1,2^, has not changed in the last two decades, and median survival is less than 14 months after diagnosis^2^. Unlike many solid tumors where immune checkpoint blockade (ICB) has demonstrated efficacy, both off-the-shelf immunotherapies^3–5^ and personalized antigen-directed therapies^6,7^ have largely failed to improve outcomes for patients with GBM, suggesting that fundamental differences exist in the tumor-immune microenvironment of brain tumors.

For nearly a century the central nervous system (CNS) was considered to be an immunologically privileged site incapable of mounting meaningful *de novo* immune responses^8,9^. However, this paradigm has been fundamentally challenged by the emerging evidence of neuroimmune communication networks, particularly at CNS border sites^10^. Despite these conceptual advances, the mechanisms governing the generation of productive anti-tumor immunity in the brain remain incompletely understood. In extracranial tumors, the mechanisms orchestrating anti-tumor immune responses are well-characterized: dendritic cells (DCs) are the antigen-presenting cells (APCs) that coordinate immunity by acquiring tumor antigens, migrating to tumor-draining lymph nodes (tdLNs) via afferent lymphatic vasculature in a CCR7-dependent manner, and cross-presenting antigens to prime CD8⁺ T cells.^11,12^ However, while DCs are enriched in CNS-adjacent tissues such as the choroid plexus and meninges^13^, the healthy brain lacks DCs and is instead populated by APC subsets including microglia and astrocytes whose roles in systemic immunity are unclear. Furthermore, the brain parenchyma lacks the network of lymphatic channels that classically mediate APC trafficking to tdLNs^10^.

RT is a cornerstone of GBM treatment and, beyond its well-established direct tumor cell killing effects, has been shown to modulate anti-tumor immune responses^14–16^. Importantly, transplant models of GBM show synergy between RT and ICB^17,18^ yet multiple clinical trials show no survival benefit^3–5^, demonstrating critical gaps in our understanding of whether and how RT shapes intracranial immunity. This is compounded in part by the treatment paradigm of resection prior to chemoradiation^2^, which limits the clinical samples available to study therapy-induced changes.

Here, we interrogate the mechanisms underlying immune dysfunction in GBM and establish a mechanistic and therapeutic framework for generating productive anti-tumor immunity in brain tumors. Using an antigen-expressing model of GBM we identify how CNS-specific DC dysfunction constrains anti-tumor immunity against GBMs. We identify microglia-mediated suppression of DC activation and migration as a critical mechanism of immune evasion and demonstrate that blocking this signaling restores endogenous anti-tumor immunity. We further demonstrate that RT reactivates DC-mediated adaptive immunity and combines with tumor antigen-targeted mRNA vaccination to drive durable tumor control mediated by systemic CD8^+^ T cells. This therapeutic approach is effective in both orthotopic GBMs and a novel genetically engineered mouse model of GBM. These findings provide a mechanistic rationale for combining RT with personalized cancer vaccines and establish DC-mediated immunity as a critical therapeutic target in brain tumors.

## Results

### GBMs drive accumulation of cDC2s, which are dispensable for anti-tumor immunity

To interrogate the dysfunctional intracranial immune responses observed in GBM, we engineered the CT2A GBM cell line^19^ to express the antigenic minimal ovalbumin (minOva) protein, containing MHC-I-restricted (SIINFEKL) and MHC-II-restricted (ISQAVHAAHAEINEAGR) epitopes, fused to the pH-stable fluorophore zsGreen, which labels tumor cells and APCs carrying tumor antigens^20^, termed CT2A-zsGreen-Ova (CzO) tumors. When we implanted CzO tumors into the flanks of syngeneic immunocompetent mice, 40% of tumors were spontaneously rejected (**Figure 1A**). However, when CzO tumor cells were implanted intracranially, tumors progressed significantly more quickly and were uniformly fatal (**Figure 1A**). These tissue-specific differences in growth of the same tumor cell line suggest that a CNS-specific mechanism of immune evasion promotes brain tumor progression. Treatment with combined anti-CTLA-4 and anti-PD-1 ICB induced complete rejection of flank tumors, but failed to control intracranial tumor growth, despite the strong antigenicity of CzO tumors (**Figure S1A**). We observed that only flank tumors induced systemic immunity, while brain tumors failed to initiate measurable systemic immune responses (**Figure 1B**). Consistent with published literature demonstrating the necessity of conventional dendritic cells (cDCs) to initiate anti-tumor immune responses^21–23^, we found that systemic immunity elicited by flank CzO tumors was abrogated in *Zbtb46^DTR^*mice after depletion of cDCs (**Figure 1C**). This was consistent with a loss of spontaneous flank tumor rejection in mice lacking type 1 conventional dendritic cells (cDC1s) (**Figure S1B**).

**Figure 1.**
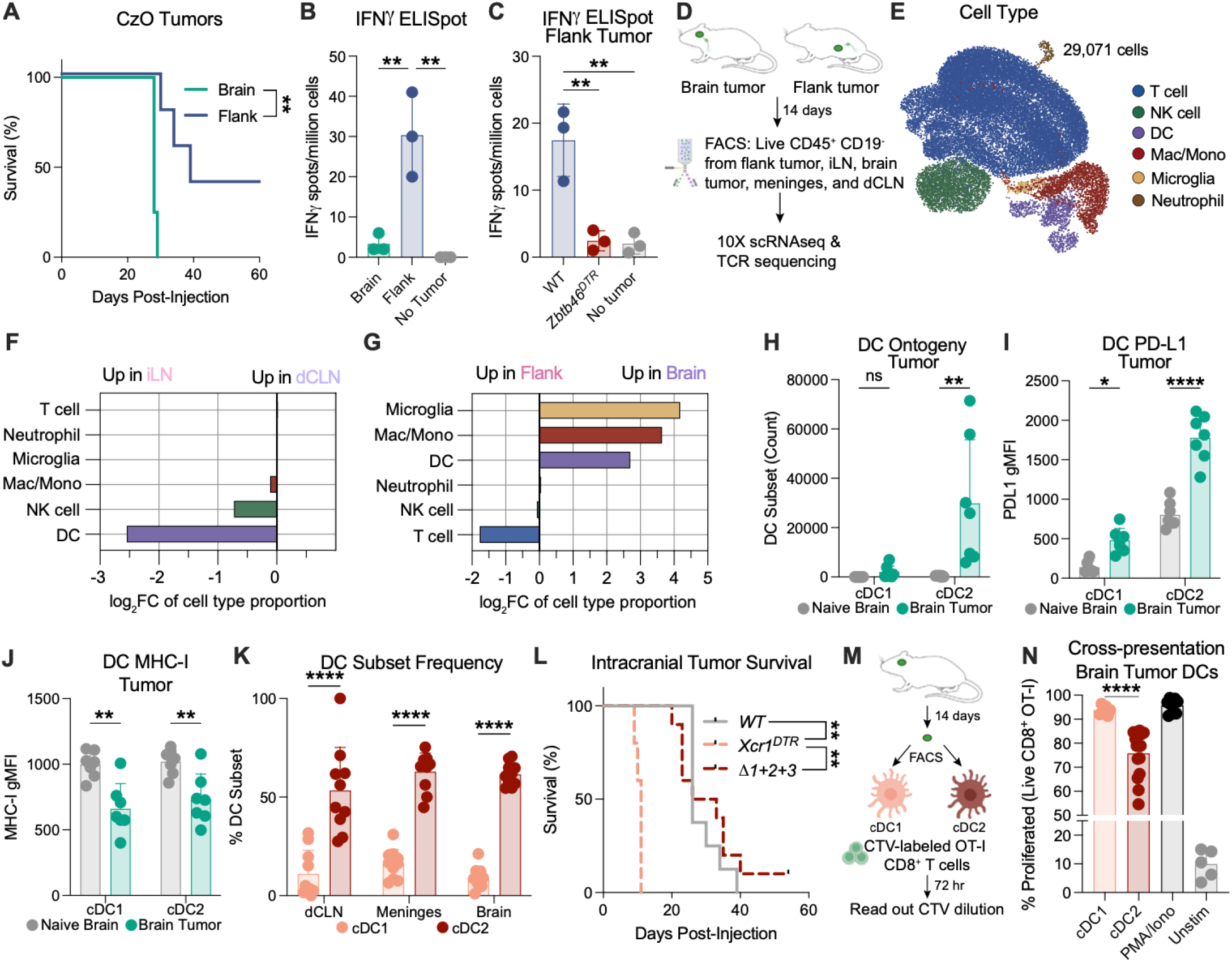
Systemic immunity depends on cDC1s, but GBMs drive accumulation of weakly antigen-presenting cDC2s. (A) Kaplan-Meier plot showing overall survival for mice with either intracranial (brain) or flank CzO tumors (n=4-5 per group; one independent experiment shown). (B) ELISpot quantification of IFNg-producing splenocytes 7 days after brain, flank, or no CzO tumor implantation (n = 3 mice per group; one independent experiment shown). (C) ELISpot quantification of IFNg-producing splenocytes 7 days after flank CzO tumors were implanted into WT chimeras, *Zbtb46^DTR^* chimeras, or no tumor control. Diphtheria toxin was injected beginning 1 day prior to tumor implantation for all groups (n = 3 mice per group; one independent experiment shown). (D) Experimental schematic for scRNA-sequencing and TCR sequencing of brain or flank tumors, deep cervical lymph node (dCLN), inguinal lymph node (iLN), and meninges. (E) UMAP plot of all 29,071 cells from brain tumor, flank tumor, dCLN, iLN, and meninges analyzed by scRNA-seq, colored by cell annotation including T cells, natural killer (NK) cells, dendritic cells (DC), macrophages/monocytes (Mac/Mono), microglia, and neutrophils. (F) Log_2_ fold change (FC) of cell type proportion between iLN and dCLN. (G) Log_2_ fold change (FC) of cell type proportion between flank and brain tumors. (H) Number of cDC1 and cDC2 in naive or GBM-bearing R hemisphere 14 days after CzO tumor injection or sham (n = 7 mice per group; one independent experiment shown). (I) Quantification of PD-L1 on cDC1 or cDC2 subsets in naive or GBM-bearing R hemisphere 14 days after CzO tumor injection or sham measured by geometric mean fluorescence intensity (gMFI) (n = 7 mice per group; one independent experiment shown). (J) Quantification of MHC-I on cDC1 or cDC2 subsets in naive or GBM-bearing R hemisphere 14 days after CzO tumor injection or sham measured by gMFI (n = 7 mice per group; one independent experiment shown). (K) Frequency of cDC1 and cDC2 (as % of all DCs) in dCLN, meninges, and tumor-bearing brain (n = 10 mice per group; two independent experiments). (L) Kaplan-Meier plot showing overall survival for WT, *Xcr1^DTR^*, or *Δ1+2+3* mice with intracranial CzO tumors. WT and *Xcr1^DTR^* mice were treated with diphtheria toxin beginning 1 day prior to tumor implantation (n = 5-10 mice per group; one independent experiment shown). (M) Experimental design for (N). CzO tumors were harvested 14d after implantation. cDC1 and cDC2 were isolated via FACS and co-cultured with Cell-Trace Violet (CTV)-labeled OT-I T cells. CTV dye dilution was analyzed by flow cytometry 72 hours later to assess proliferation. (N) Quantification of OT-I T cell proliferation after being cultured with cDC1s sorted from CzO tumors, cDC2s sorted from CzO tumors, PMA and ionomycin (PMA/Iono), or alone (Unstim) (10 tumors pooled for sorting; two independent experiments). Data are shown as mean ± SD. **p<0.005, ****p<0.0001; Log-rank test (A) with Holm-Šídák multiple comparison test (L), one-way ANOVA with Tukey’s multiple comparison test (B, C, N), or two-way ANOVA with Tukey’s multiple comparison test (H-K).

To define tissue-specific differences between CzO tumor immune microenvironments, we performed single-cell RNA sequencing (scRNA-seq) and T cell receptor (TCR) sequencing on live CD45^+^ CD19^−^ immune cells from three compartments: tumors (brain or flank), tdLNs (including the deep cervical lymph node (dCLN) for brain tumors and the inguinal lymph node (iLN) for the flank tumors)^24^, as well as the meninges (**Figure 1D**), which have been shown to be a secondary site for CNS immune responses^25–28^. As expected, the vast majority of both tdLNs were composed of T cells (**Figure 1E-F, S1C-E**). Flank tumors showed robust T cell and NK cell infiltration (**Figure 1G, S1D-E**), indicating intact adaptive immune responses. In contrast, brain tumors showed a unique infiltration of myeloid cells, consistent with the finding that myeloid cells are the most prevalent non-malignant cell type in human gliomas (**Figure 1G, S1E**) ^29,30^.

Although brain tumors failed to mount DC-mediated systemic immunity (**Figure 1B, C**), a significant enrichment of DCs was identified in both the meninges and brain of GBM-bearing mice (**Figure 1G, S1E**). DCs are crucial in anti-tumor immunity, and distinct DC subsets differ in their ability to capture, process, and present antigens, leading to differential ability to prime and sustain tumor-specific T cells^12^. While cDC1s serve as the principal APCs that coordinate anti-tumor immunity by acquiring tumor antigens, migrating via lymphatic vessels to tumor-draining lymph nodes (tdLNs), and cross-presenting antigen to prime CD8⁺ T cells^11,12^, the role of type 2 conventional dendritic cells (cDC2s) is less clear^31–33^. They can influence both CD4⁺ and CD8⁺ T cell states based on the environmental cues they receive, and have been shown to have both pro– and anti-inflammatory properties^31,34,35^. We compared the DC composition in mice bearing intracranial CzO GBMs 14 days after tumor injection to naive control mice. We found that, while naive and tumor-bearing brains had very low numbers of infiltrating cDC1s, the presence of intracranial GBMs drove accumulation of cDC2s (**Figure 1H**). These cDC2s in GBM-bearing brains expressed high levels of PD-L1 and low levels of MHC-I (**Figure 1I, J**), suggesting they may not be able to robustly prime or restimulate T cell responses. Interestingly, cDC2s were more abundant in all three CNS-associated tissues in GBM-bearing mice: dCLN, meninges, and tumor-bearing brain (**Figure 1K**). To determine the contribution of cDC1s and cDC2s to intracranial GBM immunity, we used genetic approaches to deplete each of these subsets. We implanted CzO tumors intracranially into either wild type (WT) mice, *Xcr1^DTR^* mice in which cDC1s are depleted upon administration of diphtheria toxin^36^, or *Δ1+2+3* mice in which cDC2 development is constitutively ablated by mutations within the *Zeb2* enhancer^37^. Despite the significant enrichment of cDC2 in GBM-bearing brains, only cDC1 depletion significantly decreased survival (**Figure 1L, S1F**).

To directly test whether this was due to ineffective T cell priming by cDC2s, we directly tested the priming capacity of intratumoral DC subsets derived from intracranial GBMs. We performed an *ex vivo* T cell priming assay in which naive OT-I T cells were activated using cDC1s or cDC2s sorted from CzO tumors (**Figure 1M, N**), which revealed that tumor-derived cDC1s induced complete proliferation of OT-I CD8^+^ T cells, while tumor-derived cDC2s induced significantly less CD8^+^ T cell proliferation. Together, these results demonstrate that, while systemic immunity is mediated by cDC1s, intracranial GBMs promote accumulation of cDC2s, which fail to efficiently stimulate CD8^+^ T cells to mount effective anti-tumor immune responses.

### Tumor-specific CD8^+^ T cells are primed in the deep cervical lymph node

The immunological hallmarks of GBM include extensive local and systemic immunosuppression^38,39^, low tumor mutational burden^40^, a paucity of tumor-infiltrating lymphocytes^41–43^, and sanctuary within the blood-brain barrier^44^. To test whether T cell responses were mounted against CzO tumors, we implanted tumors and adoptively transferred antigen-specific OT-I CD8^+^ T cells (**Figure 2A**). We identified robust accumulation of antigen-specific CD8^+^ T cells within the CzO tumors (**Figure 2B**), demonstrating that T cells can be primed against GBM antigens and accumulate in the GBM microenvironment, but the location of this T cell priming remained unclear. In GBM, T cell activation has been shown to occur in tertiary lymphoid structures within the tumor itself^45^, in the overlying cranial bone marrow^46,47^, within the meninges^26^, and within the cervical lymph nodes^48^. To test whether priming occurs within the meninges, tumor, or dCLN, we adoptively transferred OT-I T cells into tumor-bearing mice and analyzed the proliferation and activation state of the OT-I T cells 36 hours later (**Figure 2C**). At this early priming time point, significant OT-I T cell accumulation was observed only within the dCLN (**Figure 2D, S2A**). Additionally, priming CTV dilution peaks were present exclusively within the dCLN, while the rare OT-I T cells that were present in the meninges and the tumor had already undergone proliferation (**Figure 2E**). Furthermore, endogenous (non-OT-I) T progenitor exhausted cells (T_pex_), the stem-like memory cells that sustain long-term anti-tumor immunity^51^, were enriched specifically in the dCLN (**Figure S2B**).

**Figure 2.**
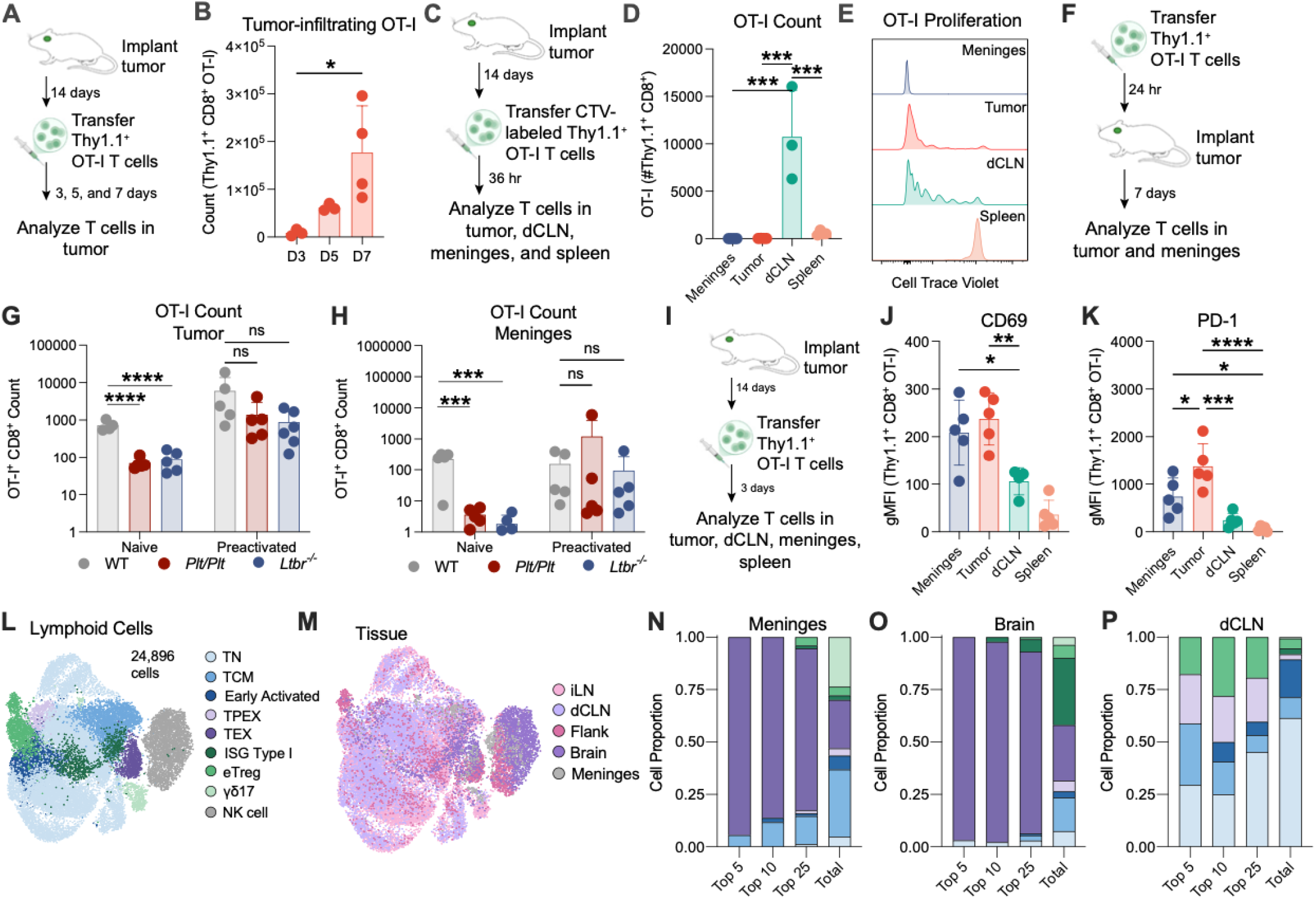
GBM-specific CD8^+^ T cell responses are primed in the dCLN. (A) Experimental design for (B). 14 days after CzO tumor implantation, 5×10^5^ OT-I CD8^+^ T cells were adoptively transferred and tumors were harvested 3, 5, or 7 days later. (B) Quantification of OT-I CD8^+^ T cells in GBM-bearing R hemisphere (n = 3-4 mice per group; one independent experiment shown). (C) Experimental design for (C). 14 days after CzO tumor implantation, 5×10^5^ CTV-labeled OT-I CD8^+^ T cells were adoptively transferred and tumors, dCLN, meninges, and spleen were harvested 36 hours later. (D) Quantification of OT-I CD8^+^ T cells in indicated tissues (n = 3-4 mice per group; one independent experiment shown). (E) Representative histograms showing Cell Trace Violet dilution in OT-I CD8^+^ T cells from specified tissues (n = 3-4 mice/group; two independent experiments). (F) Experimental design for (G-H). 1 day before CzO tumor implantation, 5×10^5^ OT-I CD8^+^ T cells were adoptively transferred into WT, *Plt/Plt*, or *Ltbr^−/−^* mice and tumors and meninges were harvested 7 days later. (G) Quantification of naive or pre-activated OT-I CD8^+^ T cells in GBM-bearing R hemisphere of WT, *Plt/Plt*, or *Ltbr^−/−^* mice (n = 4-6 mice per group; one independent experiment shown). (H) Quantification of naive or pre-activated OT-I CD8^+^ T cells in meninges of WT, *Plt/Plt*, or *Ltbr^−/−^* mice (n = 5 mice per group; one independent experiment shown). (I) Experimental design for (J) and (K). 14 days after CzO tumor implantation, 5×10^5^ CTV-labeled OT-I CD8^+^ T cells were adoptively transferred and tumors, dCLN, meninges, and spleen were harvested 3 days later. (J) Quantification of CD69 on OT-I CD8+ T cells from meninges, tumor, dCLN, or spleen measured by geometric mean fluorescence intensity (gMFI) (n = 4-5 mice per group; one independent experiment shown). (K) Quantification of PD-1 on OT-I CD8+ T cells from meninges, tumor, dCLN, or spleen measured by geometric mean fluorescence intensity (gMFI) (n = 5 mice per group; one independent experiment shown). (L) UMAP plot of all 24,896 lymphoid cells from brain tumor, flank tumor, dCLN, iLN, and meninges analyzed by scRNA-seq, colored by cell annotation including naive T cells (TN), central memory T cells (TCM), early activated T cells, T progenitor exhausted cells (TPEX), exhausted T cells (TEX), T cells with type I interferon-stimulated gene program (ISG Type I), exhausted regulatory T cells (eTreg), gamma-delta 17 cells (γδ17), and natural killer (NK) cells. (M) UMAP plot of all lymphoid cells analyzed by scRNA-seq, colored by tissue of origin including inguinal lymph node (iLN), deep cervical lymph node (dCLN), flank tumor (Flank), brain tumor (Brain), and meninges. (N) Proportion of each lymphoid cell type (annotated in 2L) within the top 5, 10, 25 expanded T cell clones, or within all (total) T cells sequenced from meninges. (O) Proportion of each lymphoid cell type (annotated in 2L) within the top 5, 10, 25 expanded T cell clones, or within all (total) T cells sequenced from tumor-bearing brain. (P) Proportion of each lymphoid cell type (annotated in 2L) within the top 5, 10, 25 expanded T cell clones, or within all (total) T cells sequenced from dCLN. Data are shown as mean ± SD. *p<0.05, **p<0.01, ***p<0.001, ****p<0.0001; One-way ANOVA with Tukey’s multiple comparison test (B, D, G, H, J, K).

To test whether the tdLN contributes to T cell activation, we used two genetic models with disrupted lymphatic organs: *Plt/Plt* mice lack *Ccl19* and *Ccl21a*, preventing leukocyte chemotaxis to the lymph node^49^, and *Ltbr^−/−^* mice lack secondary lymphoid organs^50^. Notably, these strains have intact meningeal lymphatics (**Figure S2C**). In *Plt/Plt* and *Ltbr^−/−^*mice, naive T cells were unable to expand and accumulate in the tumor or meninges (**Figure 2F-H**). However, pre-activation of T cells enabled their proliferation and accumulation within both the tumor and meninges, albeit with a slightly reduced level of accumulation (**Figure 2F-H**), demonstrating that leukocyte migration to the lymph node plays a key role in T cell activation against CzO GBMs. Three days after adoptive transfer, OT-I CD8^+^ T cells with high levels of activation (CD69), cytotoxicity (Granzyme B), and antigen experience (PD-1) markers accumulated in both the meninges and the tumor (**Figure 2I-K, S2D**). These findings demonstrate that the dCLN is the site of antigen-specific CD8+ T cell priming, and that these primed T cells then migrate to the tumor and meninges to encounter antigen.

We next compared the immunophenotype of lymphoid cells within the tumor-bearing brain, meninges, and dCLN 14 days after tumor implantation to those found in flank tumors and the iLN (**Figure 2L, 2M, S2E-G**). We found that the most abundant T cell clones within the tumor and meninges exhibited an exhaustion profile (**Figure 2N, 2O, S2G-I**), with high levels of canonical exhaustion markers including *Tox, Havcr2, Lag3,* and *Tigit*, while the most expanded T cell clones within the dCLN adopted a Tpex phenotype, as well as naive, central memory, and regulatory T cell phenotypes, with exhausted T cells representing a minority of cells (**Figure 2P**). Interestingly, approximately one-third of the T cells in the tumor-bearing brain expressed a strong type I interferon-stimulated gene (ISG) program, including *Ifit1*, *Ifit3*, *Mx1*, and *Isg15* (**Figure 2O, S2J**). This signature, which has been associated with T cell exhaustion and checkpoint resistance^52^, was not observed in immunotherapy-responsive flank tumors (**Figure S2H**). Taken together, these findings reveal a reservoir of T_pex_ in the dCLN, but exhausted and dysfunctional T cell responses within the brain tumor microenvironment and meninges.

### cDC migration from brain tumors to the tdLN is impaired, preventing T-cell mediated GBM immunity

To understand how CNS-derived antigens are presented to T cells, we compared antigen uptake and presentation by DCs within the tumor and tdLN between brain and flank tumors. Both cDC1s and cDC2s from flank CzO tumors demonstrated significantly more antigen uptake (**Figure 3A, S3A**) than cDCs from brain tumors. Interestingly, MHC-I surface antigen presentation was high in cDC1s from both brain and flank tumors, but was significantly lower in brain tumor-infiltrating cDC2s (**Figure 3B**). However, the number of DCs and their expression of activation and maturation markers, including MHC-I, MHC-II, CD86, and PD-L1, was comparable between brain and flank tumors (**Figure S3B-F**), suggesting that intratumoral DC fitness is not significantly different between tumors growing at either site. Thus, we hypothesized that the ability of DCs to migrate to the tdLN and prime T cell responses there might differ between brain and flank tumors. First, we examined DCs expressing CCR7, the receptor for CCL19 and CCL21, which is upregulated upon DC maturation and is required for DC migration to the tdLN^53,54^. Strikingly, we observed a significant enrichment of CCR7^+^ DCs within brain tumors compared to flank tumors (**Figure 3C**), while CCR7^+^ DCs accumulated within the flank tdLN, the iLN, compared to the brain tdLN, the dCLN (**Figure 3D**). This was associated with a significant increase in the number of both cDC1 and cDC2 within the tdLN (**Figure 3E**), suggesting that CCR7^+^ DCs migrate from flank tumors to the iLN, but that CCR7^+^ DC migration from brain tumors to the dCLN is significantly impaired. Furthermore, DCs in the iLN displayed increased levels of MHC-I and MHC-II (**Figure 3F, 3G**), consistent with a greater degree of maturation.

**Figure 3.**
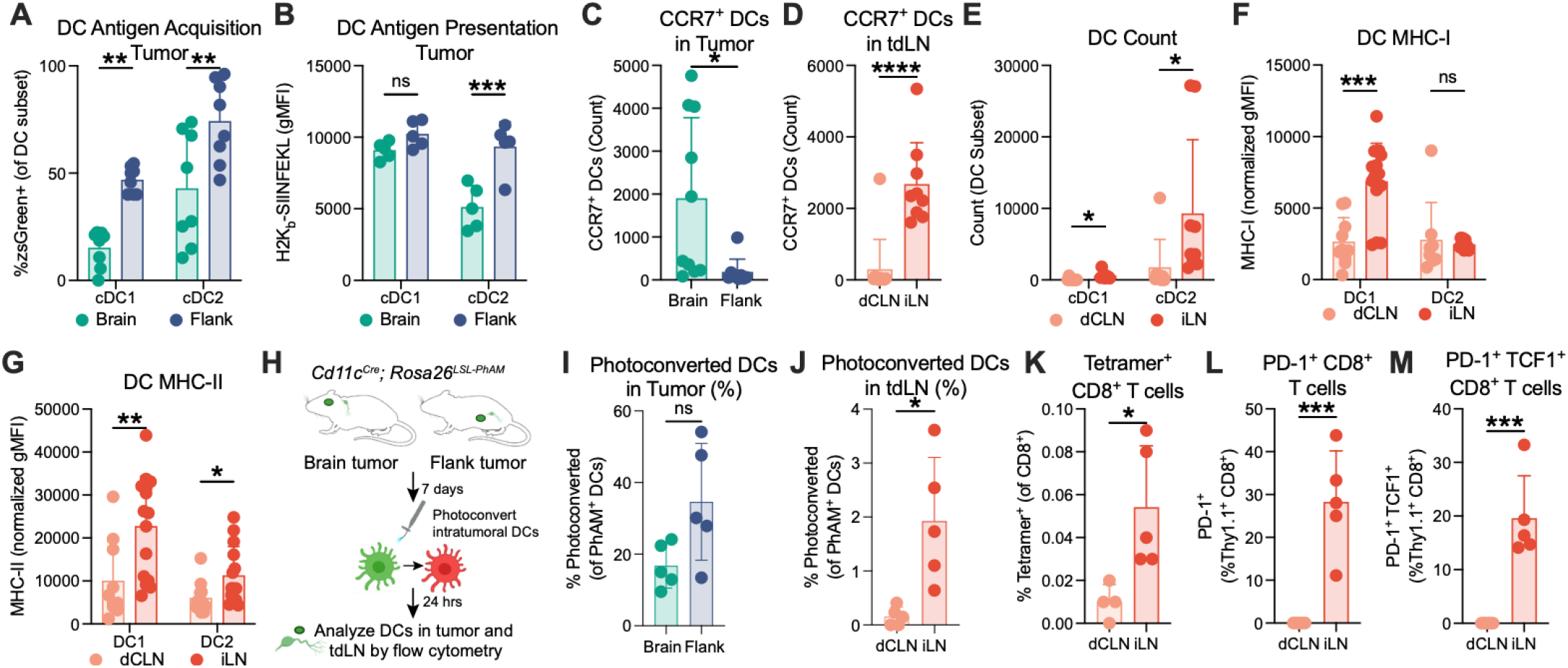
cDC migration for brain tumors to the tdLN is impaired, suppressing T cell-mediated immune responses. (A) Quantification of tumor-infiltrating zsGreen^+^ cDC1 and cDC2 in brain or flank tumors 7 days after tumor implantation. (n = 8 mice per group; two independent experiments shown). (B) Quantification of tumor-infiltrating H2Kb-SIINFEKL^+^ cDC1 and cDC2 in brain or flank tumors 7 days after tumor implantation measured by geometric mean fluorescence intensity (gMFI). (n = 5 mice per group; one independent experiment shown). (C) Quantification of tumor-infiltrating CCR7^+^ cDC1 and cDC2 in brain or flank tumors 7 days after tumor implantation. (n = 9-10 mice per group; two independent experiments shown). (D) Quantification of CCR7^+^ cDC1 and cDC2 in tdLN (dCLN for brain tumor-bearing mice, and iLN for flank tumor-bearing mice) 7 days after tumor implantation (n = 9-11 mice per group; two independent experiments shown). (E) Quantification of cDC1 and cDC2 in tdLN 7 days after tumor implantation (n = 9-11 mice per group; two independent experiments shown). (F) Quantification of cDC1 and cDC2 MHC-I expression in tdLN 7 days after tumor implantation measured by geometric mean fluorescence intensity (gMFI), normalized to mean for each experiment (n = 8-11 mice per group; two independent experiments shown). (G) Quantification of cDC1 and cDC2 MHC-II expression in tdLN 7 days after tumor implantation measured by geometric mean fluorescence intensity (gMFI), normalized to mean for each experiment (n = 8-12 mice per group; two independent experiments shown). (H) Experimental design for (I) and (J). 7 days after tumor implantation into *Cd11c^Cre;^ Rosa26^LSL-Pham^* mice, intratumoral DCs were photoconverted using a fiber optic UV probe. Tumors and tdLN were analyzed by flow cytometry 24 hr later. (I) Quantification of photoconverted DCs (as a % of all PhAM^+^ DCs) in brain or flank tumors 24 hours after photoconversion (n = 5 mice per group; one independent experiment shown). (J) Quantification of photoconverted DCs (as a % of all PhAM^+^ DCs) in tdLN 24 hours after photoconversion (n = 5 mice per group; one independent experiment shown). (K) Quantification of antigen-specific CD8^+^ T cells in the tdLN. One day before tumor implantation, 5×10^5^ OT-I CD8^+^ T cells were adoptively transferred, and antigen-specific CD8+ T cells were quantified in the tdLN 7 days later (n = 4-5 mice per group; one independent experiment shown). (L) Quantification of PD-1^+^ CD8^+^ OT-I T cells in the tdLN 7 days after tumor implantation (n = 4-5 mice per group; one independent experiment shown). (M) Quantification of PD-1^+^ TCF-1^+^ CD8^+^ OT-I T cells in the tdLN 7 days after tumor implantation (n = 4-5 mice per group; one independent experiment shown). Data are shown as mean ± SD. *p<0.05, **p<0.01, ***p<0.001, ****p<0.0001, ns: not significant; Student’s t-test (A-G, I-M).

To directly interrogate whether DC migration from brain tumors to the tdLN is impaired, we used the *Rosa26^LSL-PhAM^* mice, in which Cre recombinase can activate cell-specific expression of a photoconvertible mitochondrial-Dendra2 fluorescent protein^55^. We generated *Cd11c^Cre^; Rosa26^LSL-PhAM^*mice in which CD11c^+^ cells, including DCs, express a photoconvertible reporter. These cells emit green fluorescence at baseline, which is irreversibly converted to red upon exposure to UV light. To avoid confounding PhAM+ cells with tumor-derived zsGreen, we generated a CT2A cell line expressing cerulean and minOva (CcO) instead of zsGreen. Seven days after CcO tumor implantation, we delivered UV light by inserting a 500 µm fiber optic cable into brain or flank tumors to photoconvert tumor-infiltrating DCs (**Figure 3H**). While the proportion of photoconverted DCs was similar in brain and flank tumors (**Figure 3I**), we found a significant increase in photoconverted DCs in the iLN but not the dCLN (**Figure 3J**), demonstrating that migration from intracranial GBMs to the tdLN is indeed impaired.

Next, we aimed to assess the functional consequences of decreased DC migration to the dCLN by examining CD8^+^ T cell responses. We adoptively transferred OT-I CD8^+^ T cells prior to tumor implantation and harvested the tdLN 7 days after tumor initiation. We found that fewer CD8^+^ T cells were present in the dCLN, and that antigen-specific T cells were significantly reduced in the dCLN compared to the iLN (**Figure 3K, S3G-I**). Fewer antigen-specific T cells in the dCLN expressed PD-1 (**Figure 3L, S3J**), consistent with decreased antigen experience in the tdLN. Finally, while the iLN demonstrated robust expansion of PD-1^+^ TCF1^+^ stem-like CD8^+^ T cells, these cells, which are critical for maintaining T cell responses to cancer and are the primary responders to ICB^56–58^, were undetectable in the dCLN (**Figure 3M, S3K**). Together, these data suggest that orthotopic GBM fails to induce meaningful anti-tumor T cell responses due to impaired DC migration to the dCLN.

### Microglial signaling via AXL suppresses DC activation and migration in GBM

As glioma-infiltrating myeloid cells, which are the most prevalent nonmalignant cell type and comprise up to 50% of cells in a tumor, can recruit and suppress other immune cells^30,59^, we examined the myeloid cell landscape in our model systems (**Figure 4A, S4A, S4B**). Brain tumors showed a unique enrichment of two populations of ISG-expressing monocytes (*Isg25, Ifit1, Cxcl10, Oasl1*), while flank tumors were enriched for more classical macrophage populations (**Figure 4B**). Consistent with our previous data (**Figure 3**), we again identified an enriched migratory (CCR7^+^) DC population in iLN compared to dCLN, which was instead dominated by cDC2s, validating our flow cytometry findings. To unbiasedly understand whether the interactions between myeloid cells could contribute to the impaired DC-mediated T cell priming seen in brain tumors, we used CellChat^60^ to infer ligand-receptor interactions based on transcriptional data. Analysis of differentially expressed signaling networks between brain and flank tumors revealed that flank tumors demonstrated increased *Il15-Il15ra* signaling between neutrophils and DCs, a pattern consistent with the generation of productive systemic CD8^+^ T cell responses. In brain tumors, however, both ligands for the receptor tyrosine kinase AXL, *Pros1* and *Gas6*, were prominently expressed in microglia. *Axl* itself was expressed at increased levels on brain tumor-infiltrating DCs, with higher expression on cDC2s (**Figure 4C-E**).

**Figure 4.**
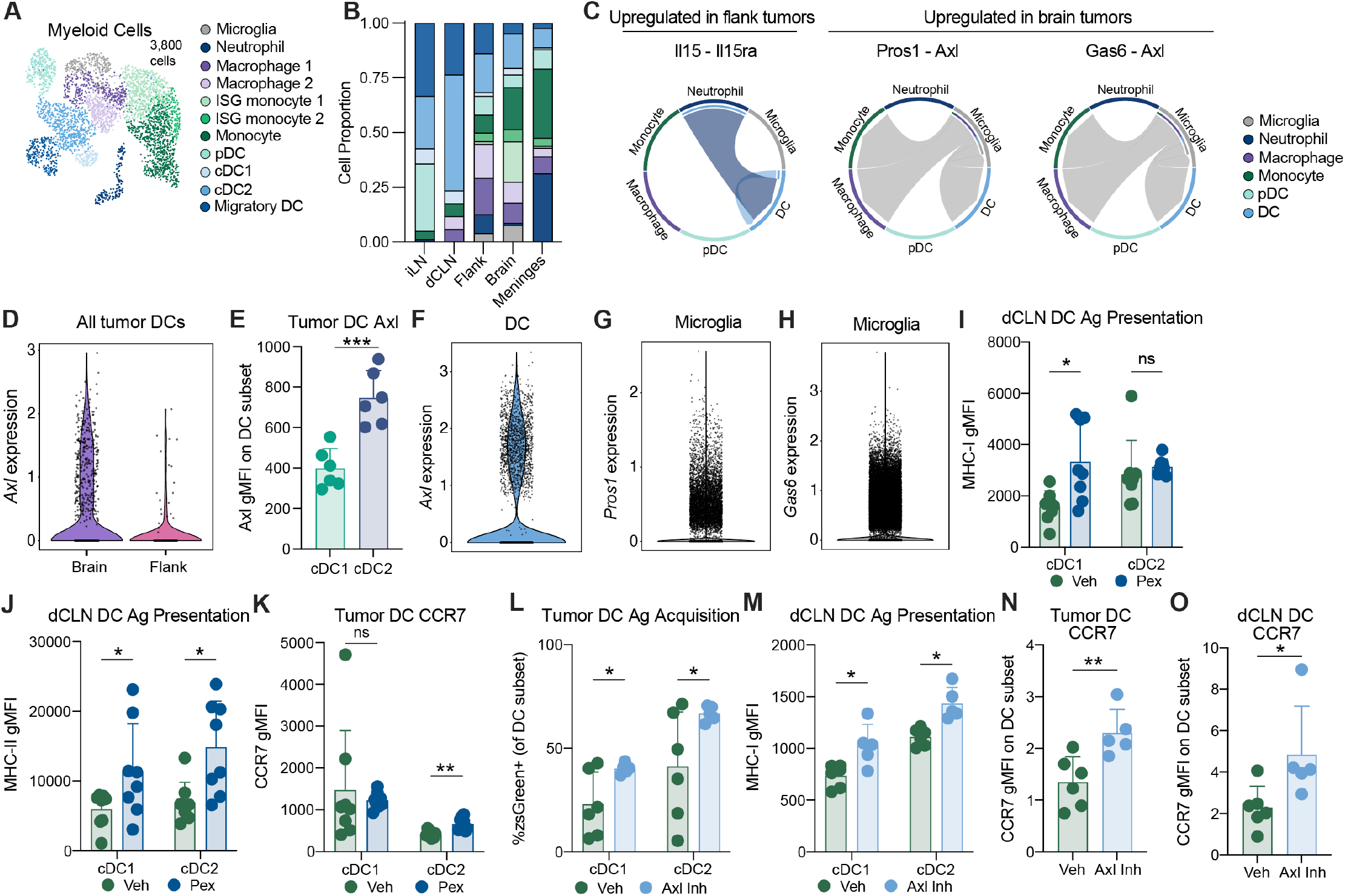
Microglial signaling via Axl suppresses DC activation and maturation in GBM. (A) UMAP plot of all 3,800 myeloid cells from brain tumor, flank tumor, dCLN, iLN, and meninges analyzed by scRNA-seq, colored by cell annotation including microglia, neutrophil, macrophage 1 and 2, interferon-stimulated gene (ISG)-expressing monocytes 1 and 2, monocytes, plasmacytoid dendritic cells (pDCs), type 1 conventional dendritic cells (cDC1), type 2 conventional dendritic cells (cDC2s), and migratory dendritic cells (Migratory DC). (B) Proportion of each cell type (annotated in 1E) within each tissue. (C) CellChat analysis of differentially expressed signaling modules demonstrating increased *Il15-Il15ra* signaling between neutrophils and DCs in flank tumors and increased *Pros1-Axl* and *Gas6-Axl* signaling between microglia and DCs in brain tumors. (D) Quantification of *Axl* expression in DCs from brain or flank tumors. (E) Quantification of *Axl* expression in cDC1s and cDC2s from brain or flank tumors 14 days after implantation (n = 6-7 mice per group; one independent experiment shown). (F) Quantification of *Axl* expression in DCs from human glioblastomas. (G) Quantification of *Pros1* expression in microglia from human glioblastomas. (H) Quantification of *Gas6* expression in microglia from human glioblastomas. (I) Quantification of cDC1 and cDC2 MHC-I expression in dCLN 14 days after tumor implantation in mice treated with vehicle (Veh) or pexidartinib (Pex) beginning 1 day prior to tumor implantation (n = 8 mice per group; one independent experiment shown). (J) Quantification of cDC1 and cDC2 MHC-II expression in dCLN 14 days after tumor implantation in mice treated with Veh or Pex beginning 1 day prior to tumor implantation (n = 8 mice per group; one independent experiment shown). (K) Quantification of cDC1 and cDC2 CCR7 expression on tumor-infiltrating DCs 14 days after tumor implantation in mice treated with vehicle (Veh) or pexidartinib (Pex) beginning 1 day prior to tumor implantation (n = 8 mice per group; one independent experiment shown). (L) Quantification of cDC1 and cDC2 zsGreen on tumor-infiltrating DCs 14 days after tumor implantation in mice treated with vehicle (Veh) or bemcentinib (Axl Inhib) beginning 1 day prior to tumor implantation (n = 6 mice per group; one independent experiment shown). (M) Quantification of cDC1 and cDC2 MHC-I on dCLN DCs 14 days after tumor implantation in mice treated with vehicle (Veh) or bemcentinib (Axl Inhib) beginning 1 day prior to tumor implantation (n = 6 mice per group; one independent experiment shown). (N) Quantification of CCR7 on tumor-infiltrating DCs 14 days after tumor implantation in mice treated with vehicle (Veh) or bemcentinib (Axl Inhib) beginning 1 day prior to tumor implantation (n = 6 mice per group; one independent experiment shown). (O) Quantification of CCR7 on dCLN DCs 14 days after tumor implantation in mice treated with vehicle (Veh) or bemcentinib (Axl Inhib) beginning 1 day prior to tumor implantation (n = 6 mice per group; one independent experiment shown). Data are shown as mean ± SD. *p<0.05, **p<0.01, ***p<0.001, ns: not significant; Student’s t-test (E, I-O).

To validate the clinical relevance of a microglial-DC AXL signaling pathway, we examined expression of *AXL* and its ligands in scRNA-sequencing from patient GBM samples^61^. In these clinical samples, *Gas6* and *Pros1* were highly expressed in microglia, and *Axl* was indeed highly expressed in GBM-infiltrating DCs (**Figure 4F-H, S4C**). In lung tumors, AXL can limit cholesterol mobilization and thus inhibit maturation of DCs^62^, so we hypothesized that AXL signaling from microglia to DCs may limit the function of brain tumor-infiltrating cDCs and constrain their ability to migrate to the tdLN.

To test whether microglia suppress cDCs, we depleted microglia using pexidartinib, a BBB-penetrant CSF1R inhibitor. We found that depletion of microglia improved antigen presentation by dCLN cDCs and increased expression of CCR7 on cDC2s in the dCLN (**Figure 4I-K**). To determine whether targeted AXL inhibition enhances the function of cDCs *in vivo*, we used the drug bemcentinib, a selective small molecule AXL inhibitor. AXL blockade resulted in significantly increased antigen acquisition by tumor-infiltrating cDCs, upregulated antigen presentation by dCLN DCs, and drove elevated CCR7 expression on both tumor-infiltrating and dCLN cDCs (**Figure 4L-O**). Together, these results demonstrate that signaling from microglia to cDCs via the AXL receptor tyrosine kinase suppresses cDC function in mouse and human brain tumors, and inhibition of AXL improves anti-tumor immunity.

### Radiation therapy increases cDC-mediated endogenous T cell immunity

Radiation therapy (RT) is a cornerstone of GBM treatment, and, in addition to its well-described direct tumor killing effects, has also been shown to modulate anti-tumor immunity^63–65^. To test whether the effects of RT were mediated by adaptive immune responses in poorly immunogenic intracranial CzO tumors, we generated tumors in WT mice or mice lacking αβ and ɣδ T cells (*TCRαβ^−/−^ɣδ^−/−^*)^66^ and treated them with either 0 Gy or 12 Gy RT. While untreated tumors did progress slightly more quickly in the absence of functional T cells, the response to high-dose single fraction RT was significantly abrogated in T cell-deficient mice (**Figure 5A**). To test whether cDCs mediated this T cell activation in response to RT, we generated bone marrow chimeras with either WT or *Zbtb46^DTR^* marrow. After bone marrow engraftment, tumors were implanted and cDCs were depleted using DT beginning 1 day prior to RT (**Figure 5B**). Mice with intact cDCs had significantly improved survival following RT (**Figure 5C**), demonstrating that DC-mediated immunity contributes to the RT response in GBM. To test which subset of cDCs contributed to RT responses in these tumors, we implanted CzO tumors intracranially into either WT, *Xcr1^DTR^*, or *Δ1+2+3* mice and found that only cDC1 depletion significantly decreased survival (**Figure 5D**). To directly test whether cDCs stimulated improved RT responses by increasing CD8^+^ T cell priming, we co-cultured either antigen-bearing or non-antigen-bearing DCs from tumors after 0 Gy or 12 Gy RT with OT-I CD8^+^ T cells (**Figure 5E**). While antigen-bearing cDCs in both conditions improved CD8^+^ T cell proliferation, we found that cDCs from irradiated tumors were able to stimulate significantly greater T cell proliferation (**Figure 5F**). To identify the mechanism by which cDCs improved CD8^+^ T cell priming after RT, we examined previously published markers of improved cDC maturation and stimulatory potential following RT^14,16,67,68^, and found no RT-induced differences in cDC1 or cDC2 abundance, frequency, antigen uptake, activation, or PD-L1 expression (**Figure S5A-L**).

**Figure 5.**
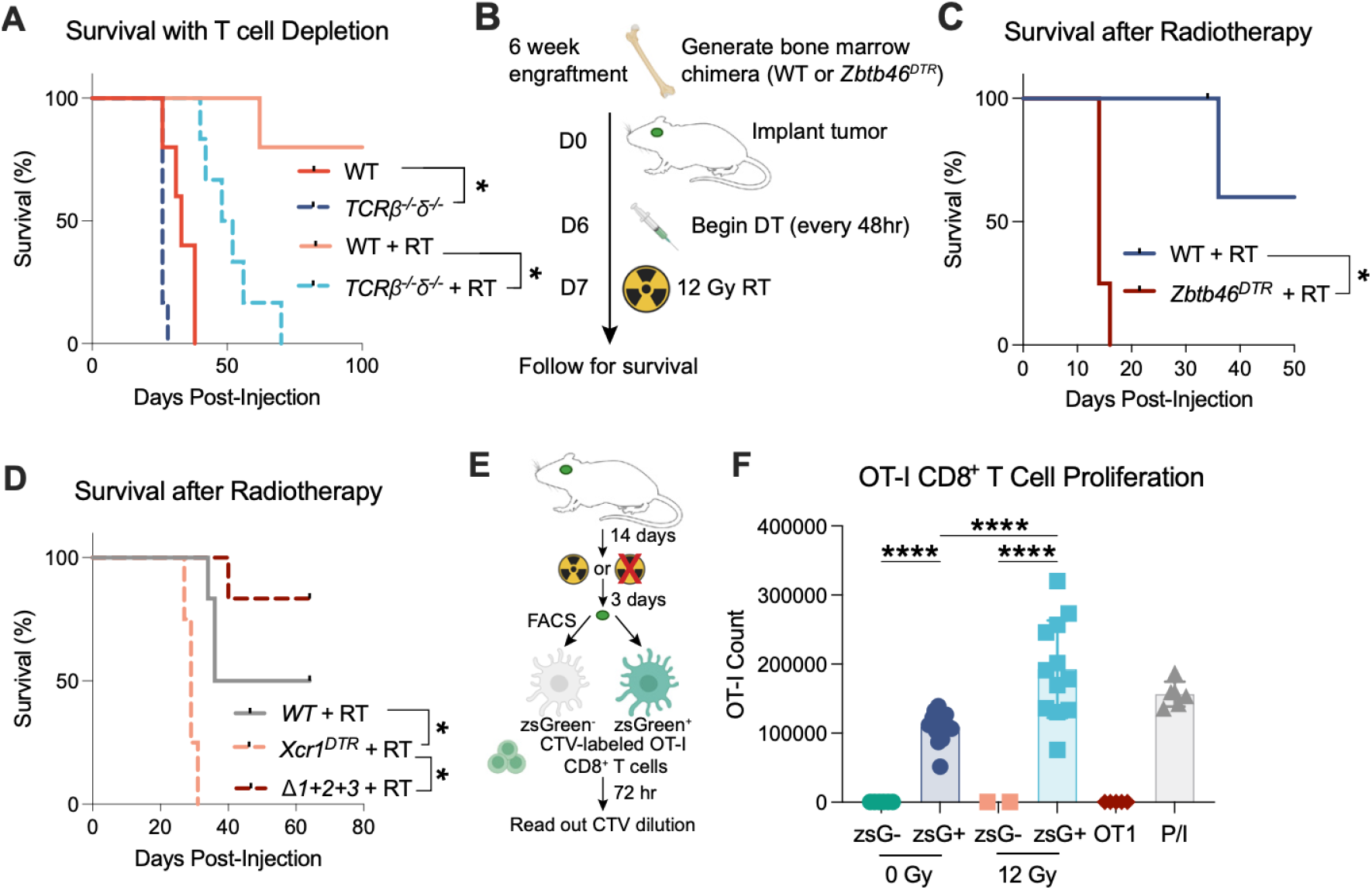
Radiation therapy activates cDC1-mediated adaptive immunity. (A) Kaplan-Meier plot showing overall survival for WT or *TCRαβ^−/−^ɣδ^−/−^* mice treated with either 0 Gy or 12 Gy RT on D7 after CzO tumor implantation (n = 5-6 mice per group; one independent experiment shown). (B) Experimental design for (C). Mice received total body irradiation followed by bone marrow transplant with either WT or *Zbtb46^DTR^* bone marrow. 6 weeks later, CzO tumors were implanted intracranially. Diphtheria toxin was injected 6 days after tumor implantation and continued every other day until endpoint. Radiation (12 Gy) or sham RT was delivered on D7 after tumor implantation and mice were followed for survival. (C) Kaplan-Meier plot showing overall survival for WT or *Zbtb46^DTR^* chimeras treated with DT delivered beginning D6 after intracranial CzO tumor implantation and either 0 Gy or 12 Gy RT on D7 (n = 4-10 mice per group; one independent experiment shown). (D) Kaplan-Meier plot showing overall survival for WT, *Xcr1^DTR^*, or *Δ1+2+3* mice with intracranial CzO tumors after RT (12 Gy) on D7. Diphtheria toxin was injected 6 days after tumor implantation and continued every other day until endpoint (n = 4-6 mice per group; one independent experiment shown). (E) Experimental design for (F). Mice with CzO tumors were treated with RT (12 Gy) or sham 14 days after implantation. zsGreen^+^ and zsGreen^−^ DCs were isolated via FACS 3 days later and co-cultured with Cell-Trace Violet (CTV)-labeled OT-I T cells. CTV dye dilution was analyzed by flow cytometry 72 hours later to assess proliferation. (F) Quantification of OT-I T cell proliferation after being cultured with DCs sorted from irradiated or unirradiated CzO tumors, PMA and ionomycin (PMA/Iono), or alone (Unstim) (10 tumors pooled for sorting; two independent experiments). Data are shown as mean ± SD. *p<0.05; ****p<0.0001. Log-rank test (C) with Holm-Šídák multiple comparison test (A, D); One-way ANOVA with Tukey’s multiple comparison test (F).

### Radiation therapy and mRNA vaccination cooperate to recruit peripheral immunity to cure orthotopic GBMs

Lipid nanoparticle (LNP)-encapsulated mRNA vaccines can generate robust antigen-specific T cell immunity against infectious agents and cancer^69–71^. To test whether mRNA vaccination could generate systemic immune responses that patrolled the CNS, we transferred OT-I CD8^+^ T cells then delivered a single dose of LNP mRNA vaccine encoding minOva (LNP-Ova) into the quadriceps muscle of naive mice (**Figure 6A**). In the absence of any BBB disruption, antigen-specific CD8^+^ T cells expanded significantly and could be identified in the dCLN, meninges, and brain (**Figure 6B, Figure S6A**). To test whether this vaccine-mediated immunity could be effective against GBM, we treated mice with CzO tumors with 0 or 12 Gy RT with or without LNP-Ova (**Figure 6C**). LNP-Ova alone did not significantly improve survival but RT, as previously shown, led to a significant tumor growth delay. Strikingly, combination treatment with LNP-Ova and RT led to cures in 100% of mice tested (**Figure 6D**). By MRI, we found that RT + LNP-Ova led to gradual tumor regression (**Figure 6E, Figure S6B**). To identify which immune cell subsets were enriched after RT + LNP-Ova, we performed flow cytometry, which revealed that combination therapy led to a significant enrichment in the frequency of tumor-infiltrating cytotoxic Granzyme B^+^ CD8^+^ antigen-specific T cells, as well as an increase in in the frequency of TCF1^+^ PD-1^+^ stem-like CD8^+^ T cells in the dCLN (**Figure 6F, 6G**). These data suggest that combination therapy with mRNA vaccines and RT reinvigorate CD8^+^ T cells in the tumor microenvironment and expand T_pex_ cells within the tdLN. To determine which cells mediated tumor cure by RT + LNP, we depleted CD4^+^, CD8^+^, or both CD4^+^ and CD8^+^ T cells beginning 1 day prior to RT. Depletion of either CD8^+^ T cells alone or CD4^+^ and CD8^+^ T cells, but not CD4^+^ T cells alone, led to a significant reduction in overall survival, demonstrating that responses to RT + LNP-Ova are mediated by CD8^+^ T cells (**Figure 6H**).

**Figure 6.**
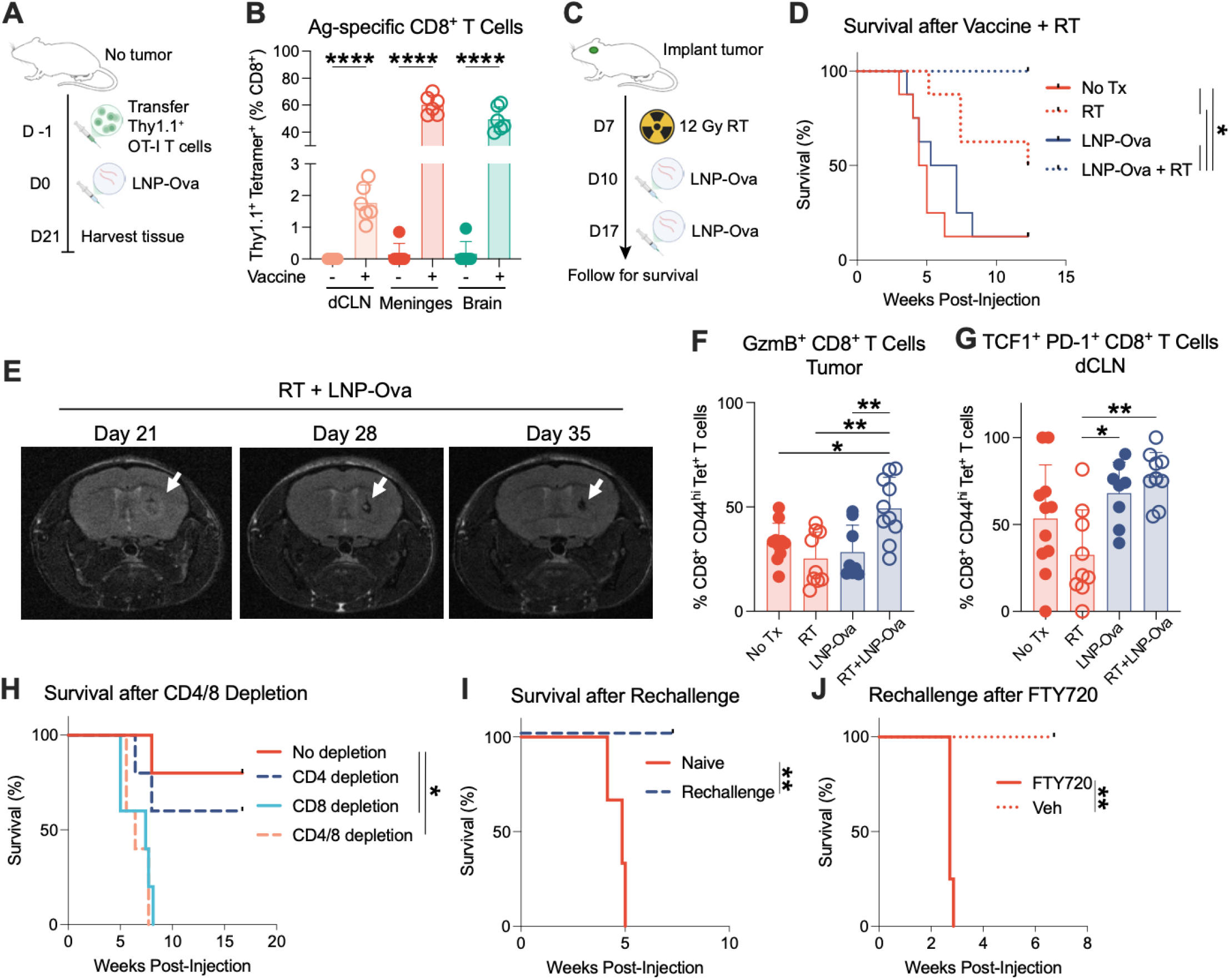
Radiation therapy and mRNA vaccination recruit peripheral immunity to cure orthotopic GBMs. (A) Experimental design for (B). Fifty thousand OT-I CD8^+^ T cells were adoptively transferred into WT mice. One day later, mice received a single dose of LNP-Ova in the quadriceps muscle. Tissue was harvested 21 days later. (B) Quantification of OT-I^+^ CD8^+^ tetramer-reactive T cells in indicated tissues 21 days after intramuscular LNP-Ova vaccination (n = 5-6 mice per group; one independent experiment shown). (C) Experimental design for (D). Intracranial CzO tumors were implanted and 7 days later, mice received 12 Gy or sham RT. On D10 and D17 after tumor implantation, mice received intramuscular LNP-Ova doses were followed for survival. (D) Kaplan-Meier plot showing overall survival for mice bearing intracranial CzO tumors treated with either no treatment (No Tx), radiation (RT, 12 Gy), LNP-Ova, or LNP-Ova and RT (n = 8 mice per group; 2 independent experiments shown). (E) MRI of CzO tumor-bearing mouse treated with RT + LNP-Ova as indicated in (C) at indicated time points after tumor implantation. Arrow indicates tumor (or prior tumor site). (F) Quantification of Granzyme B^+^ (GmzB) CD8^+^ T cells in CzO tumors treated as indicated in (C) and collected 21 days after tumor implantation. (G) Quantification of TCF1^+^ CD8^+^ T cells in the dCLN of mice with CzO tumors treated as indicated in (C) and collected 21 days after tumor implantation. (H) Kaplan-Meier plot showing overall survival for mice treated as indicated in (C) with the addition of CD4, CD8, CD4/CD8, or no depletion beginning 6 days after tumor implantation (n = 5 mice per group; one independent experiment shown). (I) Kaplan-Meier plot showing overall survival for naive mice or mice cured for 100 days after treatment with RT + LNP-Ova with subsequent CzO tumor rechallenge in the contralateral hemisphere (n = 6 mice per group; one independent experiment shown). (J) Kaplan-Meier plot showing overall survival mice cured for 100 days after treatment with RT + LNP-Ova after tumor rechallenge. One day prior to CzO tumor rechallenge in the contralateral hemisphere, mice were treated with FTY720 or vehicle (n = 5 mice per group; one independent experiment shown). Data are shown as mean ± SD. *p<0.05; **p<0.01; ***p<0.001; ****p<0.0001. One-way ANOVA with Tukey’s multiple comparison test (B, F, G). Log-rank test (I, J) with Holm-Šídák multiple comparison test (D, H).

Next, we tested whether long-term survivors had developed durable anti-tumor immunity. All surviving animals treated with RT + LNP-Ova (**Figure 6D**) were rechallenged 100 days after initial tumor implantation by CzO tumor implantation into the contralateral hemisphere and observed for survival for 8 weeks. All five naive animals died due to tumor progression by 5 weeks post-injection. In contrast, none of the long-term survivors from the RT + LNP-Ova group developed tumors, indicating that these animals had developed durable immunological memory capable of rejecting CzO tumors (**Figure 6I**). To test whether this immunological memory was mediated by CNS-resident or circulating cells, we treated mice cured by RT + LNP-Ova with FTY720 to block activated lymphocyte trafficking to tumor tissues beginning 1 day prior to tumor rechallenge and found that blockade of lymphocyte egress abrogated immunity to tumor rechallenge after combination therapy (**Figure 6J**). Taken together, these results demonstrate that RT + LNP-Ova combination treatment induces immune responses against GBM mediated by systemic circulating, rather than brain-resident, CD8^+^ T cells.

### Creation of a novel genetically engineered mouse model (GEMM) of antigen-expressing GBM

Prior studies of brain tumor immune responses are largely limited to orthotopic tumor models^42,72–75^, which do not recapitulate early interactions between evolving tumors and the immune system^76–78^. Modeling immunity during gliomagenesis is essential, as early tumor-immune interactions shape late anti-tumor immunity^79–81^, and because the treatment paradigm of resection prior to chemoradiation limits clinical samples to study therapy-induced changes. Furthermore, transplant models of GBM show synergy between RT and ICB^17,18^ yet multiple clinical trials show no survival benefit with this same therapeutic combination^3–5^. We hypothesized that mutations in PTEN and TP53, the most commonly mutated genes in GBM^82^, could be used to create a GEMM of GBM that enables tracing of the anti-tumor immune response. Mice with *Pten* and *Trp53* flanked by loxP sites (*Pten^fl/fl^; Trp53^fl/fl^* mice) were injected intracranially with a retrovirus expressing Cre recombinase and PDGFB. To express a tumor antigen, this virus also encodes the minimal ovalbumin (minOva) protein, containing MHC-I-restricted (SIINFEKL) and MHC-II-restricted (ISQAVHAAHAEINEAGR) epitopes. The minOva protein is fused to the pH-stable fluorophore zsGreen, which labels both tumor cells and APCs carrying tumor antigens (**Figure 7A**)^20^. Examination of hematoxylin and eosin-stained sections from PPzO tumors (Pten/p53 deleted, zsGreen^+^, Ova-presenting) were reviewed by a neuropathologist and found to exhibit high-grade cytologic features compatible with glial lineage. Tumor sections featured frequent mitoses and a predominantly solid growth pattern with areas of necrosis (**Figure 7B**) and focal parenchymal infiltration at the tumor periphery (**Figure 7C**). PPzO tumors demonstrated radiographic features of GBM, including irregular heterogeneous tumor margins, surrounding vasogenic edema, and a necrotic core (**Figure 7D**). In the absence of treatment, median survival was 33.5 days after tumor initiation (**Figure 7E**). Unlike many other GEMMs^80^, PPzO tumors retain antigen expression after tumor outgrowth (**Figure 7F**) and minOva epitopes are presented on tumor MHC, leading to specific cytotoxicity by antigen-specific T cells (**Figure 7G**). Unlike many non-CNS GEMMs, PPzO tumors retain persistent antigen expression, likely due to the brain tumor microenvironment’s impaired capacity for immune surveillance and immunoediting.

**Figure 7.**
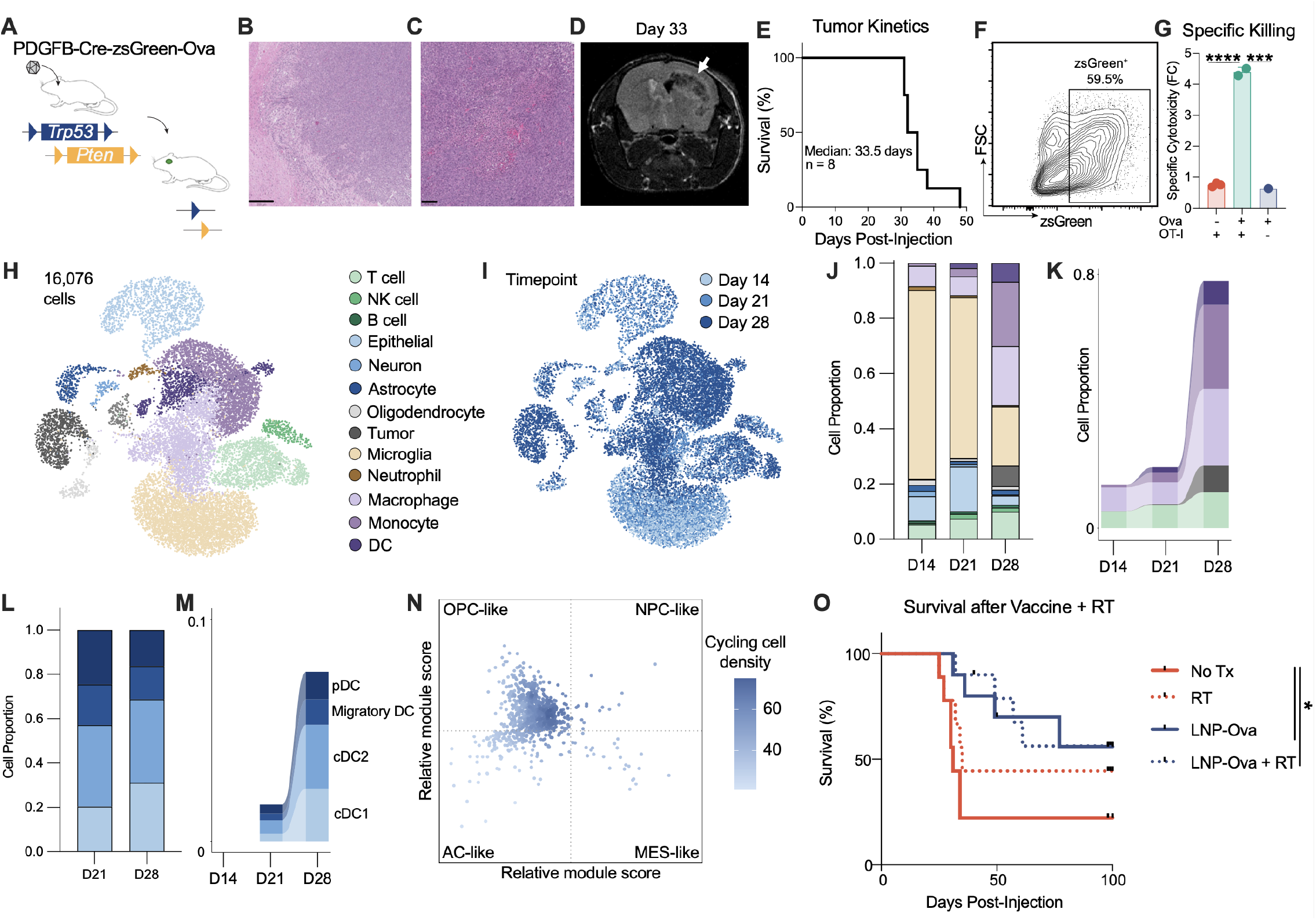
An antigen-defined autochthonous model of glioblastoma responds to mRNA vaccination and radiation therapy. (A) PPzO tumors are initiated by injection of a retrovirus encoding PDGFB, Cre, zsGreen, and minOva into *Trp53^fl/fl^; Pten^fl/fl^* mice. (B) Hematoxylin and eosin (H&E) stained sections from a representative PPzO tumor. Scale bar, 250 µm. (C) H&E stained sections from a representative PPzO tumor. Scale bar, 100 µm. (D) Representative MRI image from a PPzO tumor-bearing mouse. Arrow indicates tumor. (E) Kaplan-Meier plot showing overall survival for mice with untreated PPzO tumors. Median survival = 33.5 days (n = 8 mice). (F) Representative scatter plot of forward scatter (FSC) vs zsGreen for primary cells isolated from a PPzO tumor after 1 passage *in vitro*. (G) Quantification of specific cytotoxicity for Ova^+^ or Ova^−^ GEMM tumor cells co-cultured with or without OT-I T cells (n = 1-3 per group, one experiment shown). (H) UMAP plot of all 16,076 cells from PPzO tumors collected D14, D21, or D28 after tumor initiation analyzed by scRNA-seq, colored by cell annotation including T cells, natural killer (NK) cells, B cells, epithelial cells, neurons, astrocytes, oligodendrocytes, tumor cells, neutrophils, macrophages, monocytes, and dendritic cells (DC). (I) UMAP plot of all 16,076 cells from PPzO tumors analyzed by scRNA-seq, colored by date of collection. (J) Proportion of each cell type (annotated in 7H) within each tissue, by time point. (K) Proportion of each specified cell type (annotated in 7H) within each tissue, by time point. (L) Proportion of each dendritic cell subtype by time point. (M) Proportion of each specified dendritic cell type, by time point. (N) Two-dimensional representation of cellular states: neural-progenitor-like (NPC-like), oligodendrocyte-progenitor cell-like (OPC-like); astrocyte-like (AC-like), and mesenchymal-like (MES-like) states. Each quadrant corresponds to one cellular state, the exact position of malignant cells (dots) reflect their relative scores for the meta-modules, and their colors reflect the density of cycling cells. (O) Kaplan-Meier plot showing overall survival for mice with PPzO tumors treated with either no treatment (No Tx), radiation (RT, 12 Gy), LNP-Ova, or LNP-Ova and RT (n = 8-10 mice per group; 2 independent experiments shown). Data are shown as mean ± SD. *p<0.05; ***p<0.001; ****p<0.0001. One-way ANOVA with Tukey’s multiple comparison test (G). Log-rank test with Holm-Šídák multiple comparison test (O).

### RT and mRNA vaccination are active against autochthonous GBMs

To understand the immune milieu of our autochthonous PPzO GBMs, we performed scRNA-seq at 14, 21, and 28 days after tumor initiation. As expected, we observed an enrichment in *Pdgfra*-expressing tumor cells during tumor progression, and multiple myeloid cell populations, composed of macrophages, monocytes, and DCs, accumulated within tumors during progression (**Figure 7H-K, S7A, S7B**). Within the lymphoid compartment, early tumors were enriched for effector T cells and Th2 T cells, while later-stage tumors showed enrichment of ɣδ T cells and, like our CzO tumors, T cells expressing an ISG signature (**Figure S7C-F**). Within the myeloid compartment, microglia were the most dominant cell type present at early time points, likely due to the small tumor volume leading to overrepresentation of normal brain tissue. At later time points, increases in cDCs, macrophages, and classical monocytes were seen (**Figure S7G-J**), again consistent with the myeloid cell milieu of orthotopic models and human GBMs^30^. Importantly, as was seen in CzO tumors, cDC2s were significantly enriched compared to cDC1s and migratory DCs (**Figure 7L, 7M**).

Within GBMs, heterogeneous tumor cell states have been identified within and across individual tumors^83,84^, and the mesenchymal tumor transcriptional state predicts improved survival after immune checkpoint blockade^85^. While PPzO tumors did recapitulate this intratumoral heterogeneity, the majority of tumor cells were enriched for the oligodendrocyte-progenitor (OPC)-like expression module (**Figure 7N**), consistent with PDGF overexpression as a driver of tumorigenesis in this model. Despite the decreased ICB responses seen in OPC-like tumors, we tested the efficacy of RT and LNP-Ova in this model. We found that LNP-Ova with or without RT led to significant improvements in overall survival in PPzO tumors (**Figure 7O**). Thus, combination therapy with RT and mRNA vaccination against tumor antigens may be an effective therapeutic approach for these aggressive and incurable tumors.

## Discussion

In this study, we identified a novel CNS-specific mechanism of immune dysfunction in which microglial suppression of cDC migration, combined with a profound recruitment of poorly stimulatory cDC2s prevents activation of tumor-reactive CD8^+^ T cells and systemic anti-tumor immunity. Specifically, we identify the dCLN as the site of T cell priming by DCs, and we find that decreased cDC migration from the brain to the dCLN blunts endogenous anti-tumor T cell responses. We further demonstrate that microglia inhibit DC maturation and migration via AXL signaling, a mechanism that is conserved in human GBM-associated DCs.

Pharmacologic AXL inhibition restores DC activation and migration. Critically, we show that RT reactivates cDC1-mediated endogenous adaptive immunity. RT synergizes with intramuscular mRNA vaccination to achieve complete tumor regression and durable, circulating immunity capable of controlling recurrent tumors. These findings establish DC-mediated immunity as a therapeutic target in brain tumors and provide mechanistic and clinical rationale for combining radiation therapy with personalized cancer vaccines.

One surprising finding of this study is the distinct tissue-specific responses to implantation of identical tumor cells: while flank CzO tumors were spontaneously rejected and responded to ICB, intracranial CzO tumors progressed quickly and were refractory to ICB despite possessing identical tumor antigens. This discrepancy suggests that either anti-tumor immune responses are generated but remain contained within the brain, or that systemic immune responses are fundamentally not generated against antigens arising in the CNS. This distinction may explain the discordance observed between immunotherapy responses in primary brain cancers versus brain metastases^3–5,86,87^, where the initial immune response arises outside the CNS and may subsequently be recruited into the brain. Supporting this hypothesis, case reports of immunosuppressed organ recipients who developed disseminated GBM after receiving organs from donors deceased of GBM indicate that systemic immune responses against GBM *may* be generated and are capable of controlling tumors when they arise extracranially^88^, suggesting that the CNS barrier is not to immune privilege per se, but rather to the initiation of anti-tumor immunity in brain-resident tumors. Critically, overcoming the CNS barrier to induce systemic immunity by administering an mRNA vaccine intramuscularly generates tumor-specific CD8^+^ T cells that effectively infiltrated intracranial tumors and achieved durable cures when combined with radiation therapy. Thus, the resistance of GBM to immunotherapy reflects not immune privilege of the CNS, but rather tissue-specific immune suppression in the DC-mediated antigen presentation and trafficking that prevents the initiation of systemic anti-tumor responses. This mechanistic insight suggests that future immunotherapeutic strategies for GBM should prioritize approaches that bypass CNS-intrinsic DC dysfunction by priming immunity systemically.

Prior to this study, the functional characterization of DC subsets in GBM has been hindered by the technical difficulty of tracing both antigen-bearing DCs and antigen-specific T cell responses in most model systems. Furthermore, the rarity of DCs in tumors does not preclude their ability to promote immune responses, as evidenced by the sparse accumulation of cDC1s in GBM but the significant impact of their depletion on survival. One open question that remains is the mechanism of cDC2 accumulation in GBM. cDC1 and cDC2 development are controlled by distinct transcriptional programs^89^, and GBM-derived signals may actively suppress cDC1 development or recruitment. Furthermore, cDC1s and cDC2s exhibit distinct antigen presentation strategies^90,91^, and their lineage origin may determine their ability to prime antigen-specific responses within different tissue contexts. Understanding the specific factors driving this recruitment and lineage skewing is essential for developing therapeutic strategy that could rebalance the intratumoral cDC compartment toward cDC1-mediated immunity.

Given the highly suppressive immune microenvironment of GBMs, there is increasing interest in alternative strategies to enhance T cell priming, such as personalized tumor vaccines. Both peptide and nucleic acid vaccines offer a strategy to increase tumor-specific T cell responses by targeting mutations ^6,92^. A major key challenge that remains in clinical translation of this mRNA vaccine approach is the identification of tumor antigens that will drive robust systemic immunity, a key predictor of survival after vaccine therapy^6^. One possibility is that tumor antigens that are presented not only on tumor cells, but also cross-presented by DCs, may be the ideal targets for vaccination. This work also suggests that combining radiation therapy with personalized cancer vaccines may provide synergistic benefit against GBM.

### Limitations of the study

While this study identifies DC dysfunction as a critical mechanism of GBM immune evasion, several limitations warrant discussion. First, our studies employ a model antigen system (ovalbumin) that oversimplifies the heterogeneous neoantigen landscape of human GBM. Whether mechanisms identified with a single defined epitope accurately reflect responses to endogenous neoantigens with variable MHC binding and tissue distribution remains unclear. Second, while we demonstrate impaired DC migration to the dCLN and identify AXL signaling as a contributing mechanism, additional barriers (altered lymphatic function, reduced chemokine gradients, BBB effects) likely contribute, and their relative contributions remain undefined. Third, clinical translation requires the development of personalized neoantigen vaccines, and whether vaccine-primed CD8^+^ T cells synergize with RT-reactivated DCs when targeting patient-specific neoantigens, particularly in the context of GBM clonal heterogeneity and neoantigen loss, requires investigation. Finally, while our therapeutic experiments employ single-dose radiation (12 Gy) at day 7 post-initiation with vaccines given at days 10 and 17, alternative fractionation schedules, timing relative to resection and chemotherapy, and vaccine dose modifications may substantially impact outcomes and would require clinical optimization. Despite these limitations, convergent findings across orthotopic and autochthonous models suggest that this treatment regimen warrants further evaluation.

## Resource availability

### Lead Contact

Requests for further information and resources should be directed to and will be fulfilled by the lead contact, Stefani Spranger.

### Materials availability

Plasmids generated in this study have been deposited to Addgene.

### Data and code availability

The mouse sequencing data generated in this study will be available in the Sequence Read Archive (SRA) repository upon publication. Any additional information required to reanalyze the data reported in this paper is available from the lead contact upon request.

## Acknowledgments

We thank Melissa Duquette for mouse colony maintenance and laboratory support; Paul Thompson, Judy A. Texeira, and Peter Jansen for administrative support. We thank the Koch Institute’s Robert A. Swanson (1969) Biotechnology Center, specifically the Flow Cytometry Core Facility, the Hope Babette Tang (1983) Histology Core Facility, the Barbara K. Ostrom (1978) Bioinformatics and Computing Core Facility, and the Preclinical Imaging and Testing Facility, as well as the MIT BioMicroCenter and Whitehead Institute Genome Technology Core for providing core services. This work was funded by NIH grants P30-CA014051 and U54CA283114 from the National Cancer Institute, as well as funding from the ASTRO Residents/Fellows in Radiation Oncology Seed Grant, the Young Investigator Award from Conquer Cancer, the ASCO Foundation, and the Resident Seed Grant from the RSNA Research & Education Foundation. Figures were created in part with BioRender.com.

## Author Contributions

Conceptualization, AJW, ZJR, FC, SMK, TJ, FMW, SS; methodology, AJW, FC, YC, TC, SSL, SMK, FMW, SS; formal analysis, AJW, ZJR, YC, TH, CTDV, RV, VLB, SSL; investigation, AJW, ZJR, FC, YC, LGR, HT, NSA, TC, RV, CTDV, HS, TAH, GW, CWM, SSL, CK; resources, AJW, TC, LGR, SMK; writing – original draft, AJW; writing – review & editing, AJW, RV, SS; funding acquisition, AJW, FMW, SS; supervision, SS.

## Methods

### Mice

C57BL/6 wild type mice (B6, strain #000664) were purchased from Jackson Laboratories. *Batf3^−/−^, Zbtb46^DTR^*, *Δ1+2+3*, *CD11c^Cre^*, *PhAM^LSL^*, *Pten^fl/fl^* and *Trp53^fl/fl^* mice were purchased from Jackson Laboratories and bred in-house. Mice were housed in a specific pathogen-free (SPF) facility in ventilated microisolator cages at the Koch Institute at MIT with ad libitum access to standard chow and water. Housing rooms maintained a 12-hour light/dark cycle and ambient temperature of 23°C. Female mice between 6 and 12 weeks were used for all experiments with CzO tumors. Male and female mice between 6 and 12 weeks were used for all experiments with PPzO tumors. All experimental animal procedures were approved by the Committee on Animal Care (CAC/IACUC) at MIT.

### Generation of bone marrow chimeras

Irradiations were performed on a Gammacell 40 Exactor (Best Theratronics) located in the same SPF facility where the mice were housed, host mice were irradiated with 500 cGy, allowed to recover for 6 hr, and subsequently irradiated again with 550 cGy. The next day, bone marrow was harvested from the femurs and tibias of donor mice, washed and resuspended in PBS (Gibco), and 1×10^7^ cells were injected retro-orbitally into the irradiated host mice. A period of 6 weeks was allowed for engraftment prior to the start of experiments.

### Tumor cell lines and cell culture

Parental CT2A cancer cell line (RRID: CVCL_ZJ44) was a gift from Dr. Forest White (Koch Institute for Integrative Cancer Research at MIT). CT2A cells were cultured at 37°C and 5% CO_2_ in DMEM (Gibco) supplemented with 10% FBS (Bio-Techne Corporation) and 1% penicillin/streptomycin (Gibco). Cell lines were routinely tested for mycoplasma. For tumor implantation, tumor cells were harvested by trypsinization (Gibco), washed twice with 1X PBS (Gibco) and resuspended in PBS.

### Diphtheria toxin-mediated depletion

For depletion of DTR-expressing cells in *Zbtb46^DTR^* bone marrow chimeras, *Xcr1^DTR^* mice, or control mice, 500 ng of diphtheria toxin (Sigma-Aldrich) was injected intraperitoneally 1 day prior to tumor implantation and subsequently injected every other day thereafter until analysis. WT bone marrow chimeras treated with diphtheria toxin served as the controls.

### Generation of zsGreen-Ova and cerulean-Ova expressing CT2A cells

To generate a neoantigen expression vector, the pLV-EF1α-IRES-puro vector (Addgene #85132) was linearized by digestion with BamHI and EcoRI restriction enzymes (NEB). The zsGreen-minOva insert was generated using the pCAGGS-zsGreen-minOva construct (a gift from Max Krummel at UCSF), then cloned into the linearized pLV-EF1α-IRES-puro vector using the In-Fusion cloning kit (Takara Bio). The resulting pLV-EF1α-zsGreen-minOva-IRES-puro construct was amplified and sequenced for accuracy. The CT2A cell line stably expressing zsGreen-minOva was generated by lentiviral transduction of the parental tumor line with the pLV-EF1α-zsGreen-minOva-IRES-puro construct and selection with puromycin (Gibco). Expression was confirmed using flow cytometry for zsGreen-expressing cells. We replaced zsGreen in the pLV-EF1α-zsGreen-minOva-IRES-puro construct with cerulean using Gateway Cloning (NEB) to generate a pLV-EF1α-cerulean-minOva-IRES-puro construct. The parental CT2A cell line stably expressing cerulean-minOva was generated by lentiviral transduction of the parental tumor line with the pLV-EF1α-cerulean-minOva-IRES-puro construct and selection with puromycin (Gibco). Expression was confirmed using flow cytometry for cerulean-expressing cells.

### IFNγ ELISpot

ELISpot plates (EMD Millipore) were coated overnight at 4°C with anti-IFNγ (BD Biosciences). Plates were washed and blocked with DMEM (Gibco) supplemented with 10% FBS (Bio-Techne Corporation), 1% penicillin-streptomycin (Gibco), and 1X non-essential amino acids (Gibco) for 2 hr at room temperature. Spleens were harvested from mice at day 7 post-tumor implantation and mashed through a 70 mm filter to generate a single cell suspension. Red blood cells were lysed with 1 mL of ACK Lysing Buffer (Gibco) on ice for 1 min and splenocytes were washed once with chilled RPMI (Gibco). As a positive control, splenocytes were incubated with a mixture of 100 ng/mL PMA (Sigma-Aldrich) and 1 mg/mL ionomycin (Sigma-Aldrich). Following an overnight incubation at 37°C and 5% CO_2_, plates were developed using the BD Biosciences mouse IFNg ELISpot kit, in accordance with the manufacturer’s protocol.

### Tumor implantation, monitoring, and treatment

For flank tumor injections, mice were anesthetized with isoflurane and 1×10^6^ tumor cells were injected subcutaneously in the flank. For brain tumor injections, mice were anesthetized with isoflurane or ketamine/xylazine and immobilized on a stereotaxic frame (Neurostar). The skull was drilled 2mm lateral and 1 mm anterior from bregma. A 5 µL syringe (Hamilton) containing 3 µL of cell suspension (7.5×10^4^ tumor cells in PBS) or retroviral suspension (>1×10^8^ titer units per mL) was injected at 3.25 mm deep from the skull (flow rate 1 µL/min).

Flank tumors were monitored three times weekly by caliper measurements in two dimensions. For brain tumor-bearing mice, body condition and neurologic function were monitored three times weekly. Mice were euthanized with CO_2_ if moribund, exhibiting poor body condition, neurologic symptoms, or when tumor volumes reached more than 13 mm in any dimension. CTLA-4 (BioXCell, BE0164) and PD-1 (BioXCell, BE0146) antibodies were administered on D3, 6, and 9 by intraperitoneal injection of 200 µL per dose at 1 mg/mL diluted in PBS.

### Tissue processing for flow cytometry and FACS

Mice were anesthetized with ketamine and xylazine and circulating immune cells were labeled with retroorbital intravenous injection of CD45 antibody (Thermo Fisher Scientific) for 3 minutes. After euthanasia, tumors, lymph nodes, spleens, and meninges were collected and plated in a scored 24-well plate (Falcon) containing 1 mL digestion buffer per well: Collagenase IV (Worthington) 125 units/mL; 1mM HEPES (Gibco); and DNAse (Roche) 100 units/mL in DMEM (Gibco) with 10% FBS (Bio-Techne Corporation). Tissues were mechanically dissociated with a pestle (CellTreat) and enzymatically digested for 15 min at 37°C in digestion buffer then mechanically dissociated again using a new pestle, followed by a second 15 min incubation at 37°C. Tissues were filtered through a 70 µm filter (PluriSelect) and washed with RPMI to generate a single cell suspension. Myelin was removed from brain tissue using Debris Removal Solution (Miltenyi Biotec) per manufacturer’s instructions. Red blood cells were lysed with ACK Lysing Buffer (Gibco) on ice for 1 min and washed with chilled RPMI.

### Flow cytometry and FACS staining

Cells were resuspended in FACS buffer (PBS (Gibco) with 2% FBS (Bio-Techne Corporation) and 2 mM EDTA (Thermo Fisher Scientific)) containing Fixable Viability Dye eFluor 780 (Thermo Fisher Scientific) to distinguish live and dead cells and aCD16/CD32 (clone 93, BioLegend) to prevent non-specific antibody binding, and incubated for 15 min at 4°C. Cells were then washed with FACS buffer and stained for surface proteins using fluorophore-conjugated antibodies resuspended in FACS buffer at the specified dilutions for 30 min at 4°C, with the exception of CCR7, which was stained at 37°C for 30 minutes prior to surface staining. Following surface staining, cells were washed twice with FACS buffer and analyzed directly or fixed for downstream intracellular staining and/or analysis the next day. Cell fixation was achieved using the Foxp3 Transcription Factor Fixation/Permeabilization buffer (Thermo Fisher Scientific). After fixation, cells were washed twice with FACS buffer, stained for intracellular proteins using fluorophore-conjugated antibodies resuspended in FACS buffer at the specified dilutions overnight at 4°C and then washed twice with FACS buffer prior to flow analysis. To obtain absolute counts of cells, Precision Count Beads (BioLegend) were added to samples following manufacturer’s instructions. Flow cytometry sample acquisition was performed on an LSR Fortessa or FACSymphony A3 cytometer (BD Biosciences), and the collected data was analyzed using FlowJo v10.10.0 software (TreeStar). For cell sorting, the surface staining was performed as described above under sterile conditions, and cells were acquired and sorted into co-culture media (RPMI (Gibco) containing 10% FBS (Bio-Techne Corporation), 1% penicillin-streptomycin (Gibco), 1X non-essential amino acids (Gibco), 1 mM HEPES (Gibco), and 50 mM b-mercaptoethanol (Gibco)) using a FACSAria III sorter (BD Biosciences). For CD8^+^ T cell analysis, cells were pre-gated on live, singlets, 3 minute IV-label^−^, CD45^+^, MHCII^−^, NK1.1^−^, TCRb^+^, CD8a^+^ markers. For DC analysis and cell sorting, cells were pre-gated on live, singlets, 3 minute IV-label^−^, CD45^+^, CD3e^−^, CD19^−^, NK1.1^−^, Ly6C^−/low^, MHCII^+^, F4/80^−^, and CD11c^+^. As previously published^31^, cDC1s were CD103^+^ and cDC2s were CD11b^+^. To identify SIINFEKL-reactive CD8^+^ T cells, PE-conjugated SIINFEKL tetramer (NIH Tetramer Core Facility) was added prior to the surface staining step in the flow cytometry methods described above. PE-conjugated SIINFEKL tetramer was titrated to empirically determine the optimal staining concentration.

### Adoptive transfer of OT-I T cells

TCR transgenic CD8^+^ T cells were isolated from spleens and lymph nodes of naïve OT-I TCR transgenic mice using the mouse CD8a^+^ T cell isolation kit (Miltenyi Biotec) following manufacturer’s instructions. If indicated, isolated CD8^+^ T cells were washed with PBS (Gibco) and stained with Cell Trace Violet (Thermo Fisher Scientific) following the manufacturer’s instructions. 5×10^5^ CD8^+^ OT-I T cells were transferred retro-orbitally into recipient mice and tissues were collected at the indicated time points and analyzed by flow cytometry. For naive T cell transfers into WT, *Plt/Plt*, or *Ltbr^−/−^* mice, 1×10^5^ CD8^+^ OT-I T cells were transferred retro-orbitally into recipient mice. For preactivated T cell transfers into WT, *Plt/Plt*, or *Ltbr^−/−^* mice, OT-I cells were negatively selected using biotinylated antibody cocktail (Biolegend: Ter119 [1:100], B220 [1:200], MHC-II [1:200], CD11b [1:200], CD11c [1:200], CD49b [1:200], TCR γ/δ [1:200], CD4 [1:200]) prior to overnight activation with plate-bound anti-CD3 (BD 553057) and soluble anti-CD28 (BD 553295). Following activation, 500k cells were transferred.

### *Ex vivo* DC-T cell co-culture assay

To obtain specific DC subsets, cells were FACS-sorted from tumors as described above. To obtain tumor-specific CD8^+^ T cells, TCR transgenic CD8^+^ T cells were isolated from spleens and lymph nodes of naïve OT-I TCR transgenic mice using the mouse CD8a^+^ T cell isolation kit (Miltenyi Biotec), following manufacturer’s instructions. Isolated CD8^+^ T cells were washed with PBS (Gibco) and stained with Cell-Trace Violet (Thermo Fisher Scientific) following the manufacturer’s instructions. 5×10^4^ dye-labeled CD8^+^ OT-I T cells and 1×10^4^ DCs (5:1 T cell-DC ratio) were mixed and added to each well of a V-bottom tissue culture-treated 96-well plate in co-culture media (RPMI (Gibco) containing 10% FBS (Bio-Techne Corporation), 1% penicillin-streptomycin (Gibco), 1X non-essential amino acids (Gibco), 1 mM HEPES (Gibco), and 50 mM β-mercaptoethanol (Gibco)). The cells were cultured at 37°C and 5% CO_2_ for 72 hr at which point T cell proliferation was measured by dye dilution via flow cytometry as a proxy for T cell activation.

### Photoconversion of tumor-infiltrating DCs

CcO flank or brain tumors were initiated as previously described in *Cd11c^Cre^; Rosa26^LSL-PhAM^*mice. Seven days after tumor initiation, flank tumors were surgically exposed and brain tumors were accessed via craniotomy at the original burr hole site. A 500 µm fiber optic cable attached to a UV light source was placed within the center of the tumor. The UV light source was built by focusing the light from a high precision 250mW laser (CivilLaser) on to one end of the fiber optic cable (Edmund Optics 02-532) using a 4.5mm diameter x 4.5mm focal length convex lens (Edmund Optics 47-866) secured in a custom 3D printed housing. The laser was powered using a 5V power supply. Tumors were exposed to 405nm light for 10 minutes at maximal laser output power (approximately 15mW). Placement within flank tumors was confirmed by visible cerulean fluorescence and placement within brain tumors was confirmed using stereotactic coordinates. Twenty-four hours after photoconversion, tumors and tdLNs were harvested for flow cytometry and tissues were processed as described above.

### Treatment with bemcentinib and pexidartinib

Bemcentinib (1037624-75-1) and pexidartinib (1029044-16-3) were purchased from Medchem Express. Pexidartinib was administered by oral gavage once daily at 40 mg/kg/day in 100uL 60% PEG400 (Sigma Aldrich)/40% sterile water. Bemcentinib was administered by oral gavage twice daily at 25mg/kg in 100 uL PEG400 (Sigma Aldrich). Treatment began one day prior to tumor implantation and continued until the day of tissue harvest (14 days after tumor implantation).

### Paired scRNAseq and TCRseq sample preparation, library generation, and sequencing

For CzO mouse model samples, Live CD45^+^ CD19^−^ singlet cells were FACS-sorted as described above from the tumors, tdLN, or meninges of WT mice bearing CzO flank or brain tumors 14 days after tumor implantation. For PPzO mouse model samples, Live singlet cells were FACS-sorted from the right cerebral hemisphere of mice bearing PPzO tumors at 14, 21, or 28 days after tumor initiation. Sorted cells were washed twice in PBS (Gibco) and resuspended at a final concentration of 1×10^3^ cells/mL in chilled PBS (Gibco) containing 0.04% bovine serum albumin (Research Product International). The cellular suspension was submitted to the Whitehead Institute Genome Technology Core for library generation and sequencing. Briefly, single cells were encapsulated into droplets using the 10X Genomics Chromium Controller, and the cDNA library was prepared using the Chromium Single Cell 3’ Reagent Kits v3 (10X Genomics) following manufacturer’s instructions. The resultant cDNA library was sequenced by using a NovaSeq 6000 (Illumina).

### Paired scRNAseq and TCRseq data alignment, processing, and analysis

Reads were aligned using the nfcore/scrnaseq pipeline version 4.1.0 with aligner Cell Ranger multi. The 10X Genomics GRCm39 reference was used for gene expression and the 10X Genomics GRCm38 reference was used for V(D)J alignment. Sequencing and alignment quality was evaluated based on the rank plot, mean reads per cell, mapping percentage (gene expression), and percentage of cells with productive V-J spanning pair.

For scRNAseq, cells were filtered based on number of UMIs, number of genes, and percent mitochondrial UMIs. DoubletFinder was used to remove doublets using assumed doublet rates based on the number of cells loaded per sample^93^. Data were normalized using SCTransform (2,000 variable features), regressing out percent mitochondrial and ribosomal UMIs^94^. Leiden clustering was used. Data was subsetted into lymphoid and myeloid populations and low quality clusters were removed. Dot plots were generated using DotPlot() in Seurat. Differential gene expression was performed using MAST on log-transformed RNA, regressing out percent mitochondrial UMIs, percent ribosomal UMIs, and number of UMIs^95^. Gene signature scores were calculated using AUCell^96^. CellChat was used to investigate ligand receptor interactions between cell types of interest^60^. When creating CellChat objects, the number of cells were not considered (population.size = FALSE in computeCommunProb()). Differentially expressed interactions between different tissues were calculated using identifyOverExpressedGenes() and associated functions. For analysis of human GBM^61^, the Core Gbmap was downloaded and markers of interest were plotted based on cell type.

For scTCRseq, contigs were obtained from Cell Ranger multi’s filtered_contig_annotations.csv. Only productive calls were used. Alpha chains were not used. Beta chains with > 2 UMIs were kept and cells with 0 or >1 beta chain were filtered. Clonotypes were defined as having identical V gene, J gene, and CDR3 nucleotide sequence. For ambiguous V gene calls, either assigned gene was allowed to match a clonotype. Clonotypes were merged with gene expression data based on cell barcode.

For paired scRNAseq/TCRseq analysis, clonotyping was merged with gene expression data based on matching cell barcodes. After quality control and clonotyping we recovered over 4,100 T cells with productive TCRs, 75% of which had paired expression data, resulting in over 3,000 unique clones.

### mRNA lipid nanoparticle vaccine

Lipid nanoparticles (LNPs) were synthesized via a microfluidic organic–aqueous mixing method. The ionizable lipid 9-heptadecanyl 8-{(2-hydroxyethyl)[6-oxo-6-(undecyloxy)hexyl]amino}octanoate (SM-102) was purchased from BroadPharm (BP-25499). 1,2-Distearoyl-*sn*-glycero-3-phosphocholine (DSPC; 850365), cholesterol (700100), and 1,2-dimyristoyl-*rac*-glycero-3-methoxypolyethylene glycol-2000 (DMG-PEG2k; 880150) were purchased from Avanti Polar Lipids. 10 mM citrate buffer (pH 3.0; J61391-AK) was obtained from Alfa Aesar. CleanCap OVA mRNA (L-7610-1000; TriLink BioTechnologies) was prepared as the aqueous phase by dilution in a 10 mM citrate buffer.

The organic phase was prepared by dissolving SM-102, DSPC, cholesterol, and DMG-PEG2k in ethanol at a molar ratio of 50:10:38.5:1.5, respectively. The two phases were combined at an ethanol-to-aqueous volume ratio of 1:3 and a nitrogen-to-phosphate (N:P) ratio of 5:1. Each phase was loaded into a BD syringe and assembled onto a NxGen microfluidic cartridge for mixing using a NanoAssemblr Ignite instrument (Precision Nanosystems), operated at a total flow rate of 12 mL/min and a waste volume of 0 mL. The resulting LNPs were dialyzed against 20 mM Tris Acetate buffer supplemented with 8% (w/v) sucrose using 10K MWCO Slide-A-Lyzer MINI Dialysis Cassettes (Thermo Fisher Scientific) at 25°C for 4 hours to remove residual ethanol and exchange buffer. The final LNP formulation contained mRNA at a concentration of 0.1 mg/mL.

For vaccination, mice were anesthetized with isoflurane and 50 µL of the above-described LNP solution was injected into the right quadriceps muscle.

### Small animal MRI

Magnetic resonance imaging was performed *in vivo* using a 7T MRI system operated with a Bruker AV4 Neo BioSpec 70/20 USR console (Bruker BioSpin, Rheinstetten, Germany), equipped with a 114-mm, 660-mT/m actively shielded gradient and a QSN075/040 RF coil. Mice were anesthetized with 2.5% isoflurane for induction and maintained at 2–2.5% isoflurane throughout data acquisition. Body temperature and respiratory rate were continuously monitored using an SAII monitoring and gating system (Small Animal Instruments Inc., Stony Brook, NY, USA). Mouse brain images were acquired in the transverse (axial) orientation and reconstructed using Bruker ParaVision PV360 v2.0. T2-weighted images were acquired using a TurboRARE sequence with a repetition time (TR) of 2000 ms, echo time (TE) of 30 ms, three signal averages, and a RARE factor of 8. The imaging geometry consisted of a 256 × 256 acquisition matrix, 20 × 20 mm² field of view (FOV), 20 interleaved slices, and a slice thickness of 0.5 mm with no interslice gap. Images were converted to DICOM format and reviewed using ImageJ (https://imagej.nih.gov).

### FTY720 Treatment

FTY720 (Enzo Life Sciences) stock solution at 10 mg/mL in DMSO was diluted to 125 µg/mL concentration in PBS directly before administration. Mice received a dose of 25 µg FTY720 or PBS containing DMSO as control. Treatment was given every other day for the duration of the experiment.

### Histopathology of PPzO tumors

Mice were anesthetized with ketamine and xylazine and transcardially perfused with ice-cold PBS followed by 4% PFA. Tumor tissues were harvested and immediately fixed for 24 hours in 4% PFA, then tissues were processed to paraffin, embedded, sectioned, and stained with hematoxylin and eosin (H&E). H&E sections of brain tumors were reviewed by a board-certified neuropathologist (CWM).

### Specific Killing Assay

OT-I T cells were activated for 72 hours in a plate coated with αCD3/αCD28. Target cells were PPzO GEMM lines (passage >10) generated via retroviral injection of Cre, PDGF, and zsG-minOva (Ova^+^), or Cre, PDGF, and zsG (Ova^−^). Target cells were trypsinized, washed, and counted as previously described^97^. Ova^+^ cells were incubated with 5 µM CTV and Ova^−^ cells were incubated with 100 nM CTV in PBS (Gibco) at 37C for 20 minutes. Labeling reactions were quenched with DMEM and 10% FBS and cells were pelleted at 500g for 5 min. Ova^+^ and Ova^−^ cells were mixed at a 1:1 ratio and plated into round-bottom 96-well plates at 20,000 cells per well. For Ova^−^ comparison, Ova-PPzO cells were incubated with 5 µM CTV and CT2A parental cells were incubated with 100 nM CTV in PBS (Gibco) at 37C for 20 minutes. OT-I cells were then plated into wells at 10:1 or 0:1 effector:target ratios. After 12h, cells were processed for flow cytometry. Cells were gated on live, CD45^−^ singlets and specific killing was calculated with the following formula: 100 – ((CTVhi/CTVlo – with effector cells)/(CTVhi/CTVlo without effector cells) × 100.

## Figures

**Figure S1.**
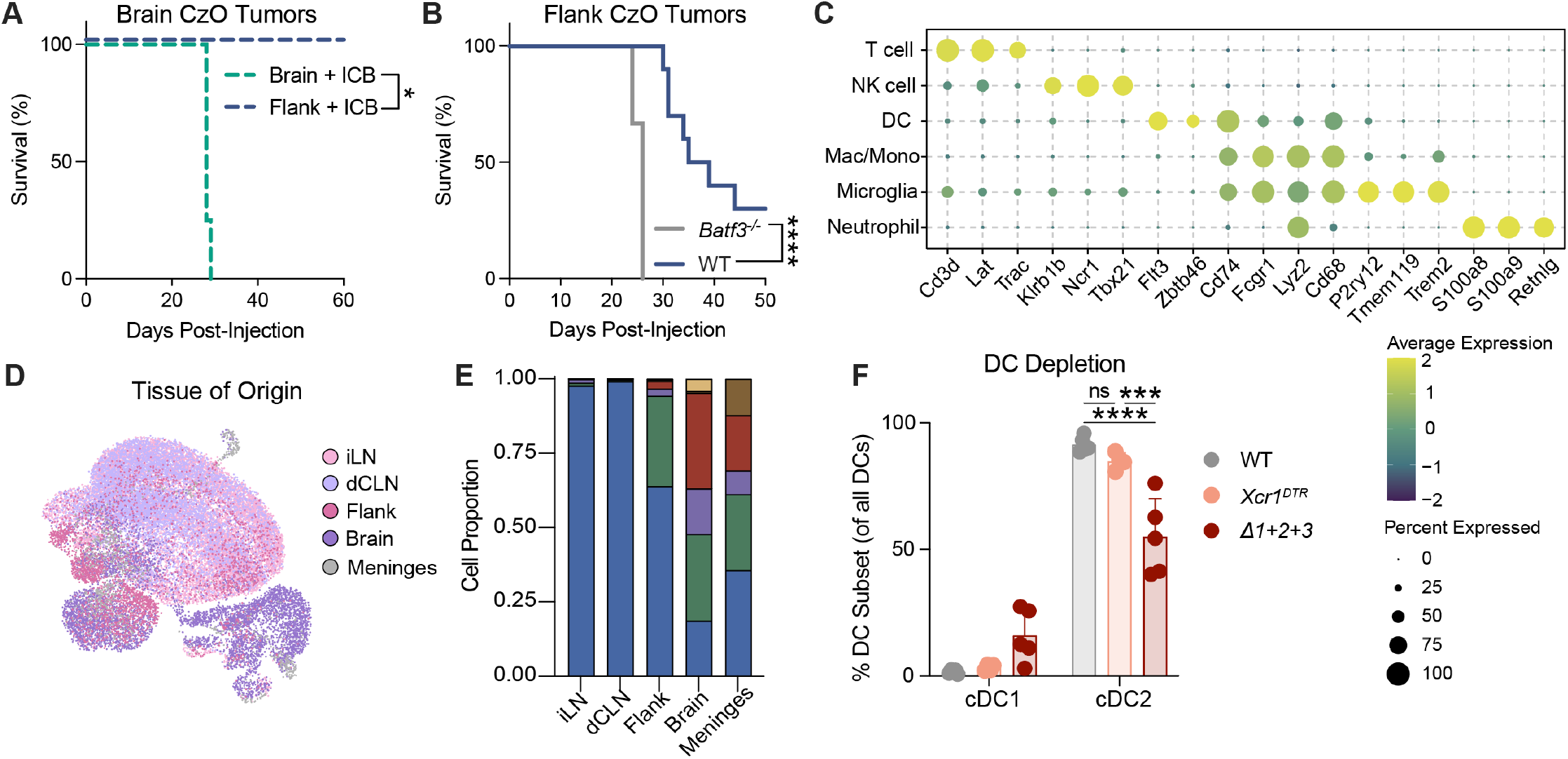
Immune responses in flank and brain CzO tumors. (A) Kaplan-Meier plot showing overall survival for mice with either intracranial (brain) or flank CzO tumors after treatment with CTLA-4 and PD-1 blockade D3, 6, and 9. (n=4-5 per group; one independent experiment shown). (B) Kaplan-Meier plot showing overall survival for WT or *Batf3^−/−^* mice lacking cDC1s with flank CzO tumors. (n=5-10 per group; one independent experiment shown). (C) Expression of subtype-defining markers. Dot size: percent expressing; color: average expression (n=29,071 cells). (D) UMAP plot of all cells analyzed by scRNA-seq, colored by tissue of origin including inguinal lymph node (iLN), deep cervical lymph node (dCLN), flank tumor (Flank), brain tumor (Brain), and meninges. (E) Proportion of each cell type (annotated in 1E) within each tissue. (F) Frequency of cDC1 and cDC2 (as % of all DCs) in tumors collected 14 days after implantation into WT, *Xcr1^DTR^*, or *Δ1+2+3* mice. (n = 4-5 mice per group; two independent experiments). Data are shown as mean ± SD. **p<0.005; ***p<0.001; ****p<0.0001. Log-rank test (A, B), two-way ANOVA with Tukey’s multiple comparison test (F).

**Figure S2.**
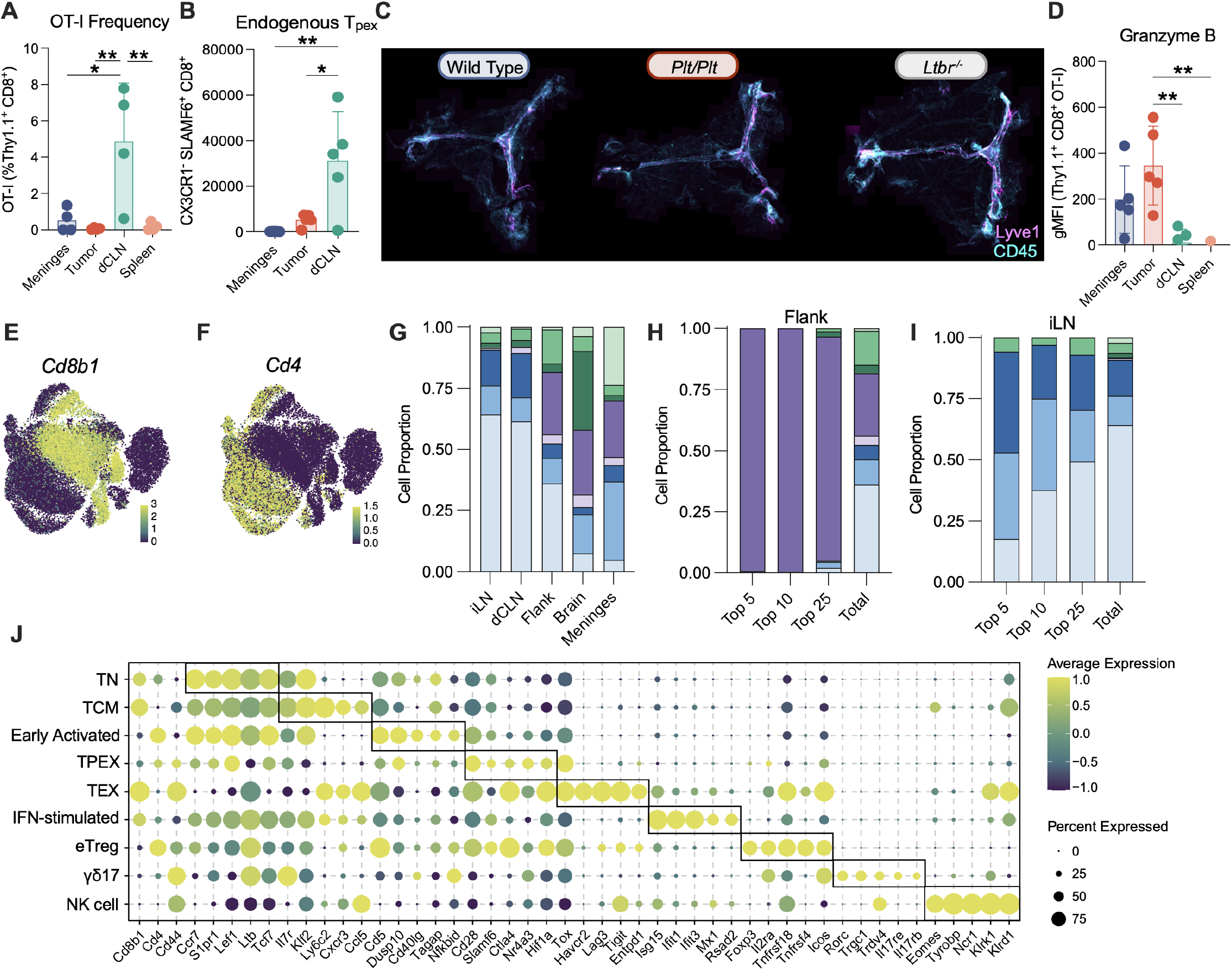
Lymphoid responses and phenotypes in CzO tumors. (A) 14 days after CzO tumor implantation, 5×10^5^ CTV-labeled OT-I CD8^+^ T cells were adoptively transferred and tumors, dCLN, meninges, and spleen were harvested 36 hours later. Quantification of OT-I CD8^+^ T cells in indicated tissues is shown (n = 4 mice per group; one independent experiment shown). (B) 14 days after CzO tumor implantation, 5×10^5^ CTV-labeled OT-I CD8^+^ T cells were adoptively transferred and tumors, dCLN, meninges, and spleen were harvested 36 hours later. Quantification of endogenous (Thy1.1^−^) CD8^+^ T cells in indicated tissues is shown (n = 5 mice per group; one independent experiment shown). (C) Immunofluorescence imaging of whole-mounted dura from WT, *Plt/Plt*, or *Ltbr^−/−^* mice. Lyve1 (pink) and CD45 (cyan) demonstrate intact meningeal lymphatics. (D) 14 days after CzO tumor implantation, 5×10^5^ CTV-labeled OT-I CD8^+^ T cells were adoptively transferred and tumors, dCLN, meninges, and spleen were harvested 3 days later. Quantification of Granzyme B on OT-I CD8^+^ T cells from meninges, tumor, dCLN, or spleen measured by geometric mean fluorescence intensity (gMFI) (n = 5 mice per group; one independent experiment shown). (E) UMAP plot of all lymphoid cells analyzed by scRNA-seq, colored by expression of *Cd8b1* transcript. (F) UMAP plot of all lymphoid cells analyzed by scRNA-seq, colored by expression of *Cd4* transcript. (G) Proportion of each cell type (annotated in 2L) within each tissue. (H) Proportion of each lymphoid cell type (annotated in 2L) within the top 5, 10, 25 expanded T cell clones, or within all (total) T cells sequenced from flank tumors. (I) Proportion of each lymphoid cell type (annotated in 2L) within the top 5, 10, 25 expanded T cell clones, or within all (total) T cells sequenced from inguinal lymph nodes. (J) Expression of subtype-defining markers. Dot size: percent expressing; color: average expression (n=24,896 cells). Data are shown as mean ± SD. *p<0.05, **p<0.01, One-way ANOVA with Tukey’s multiple comparison test (A, B, D).

**Figure S3.**
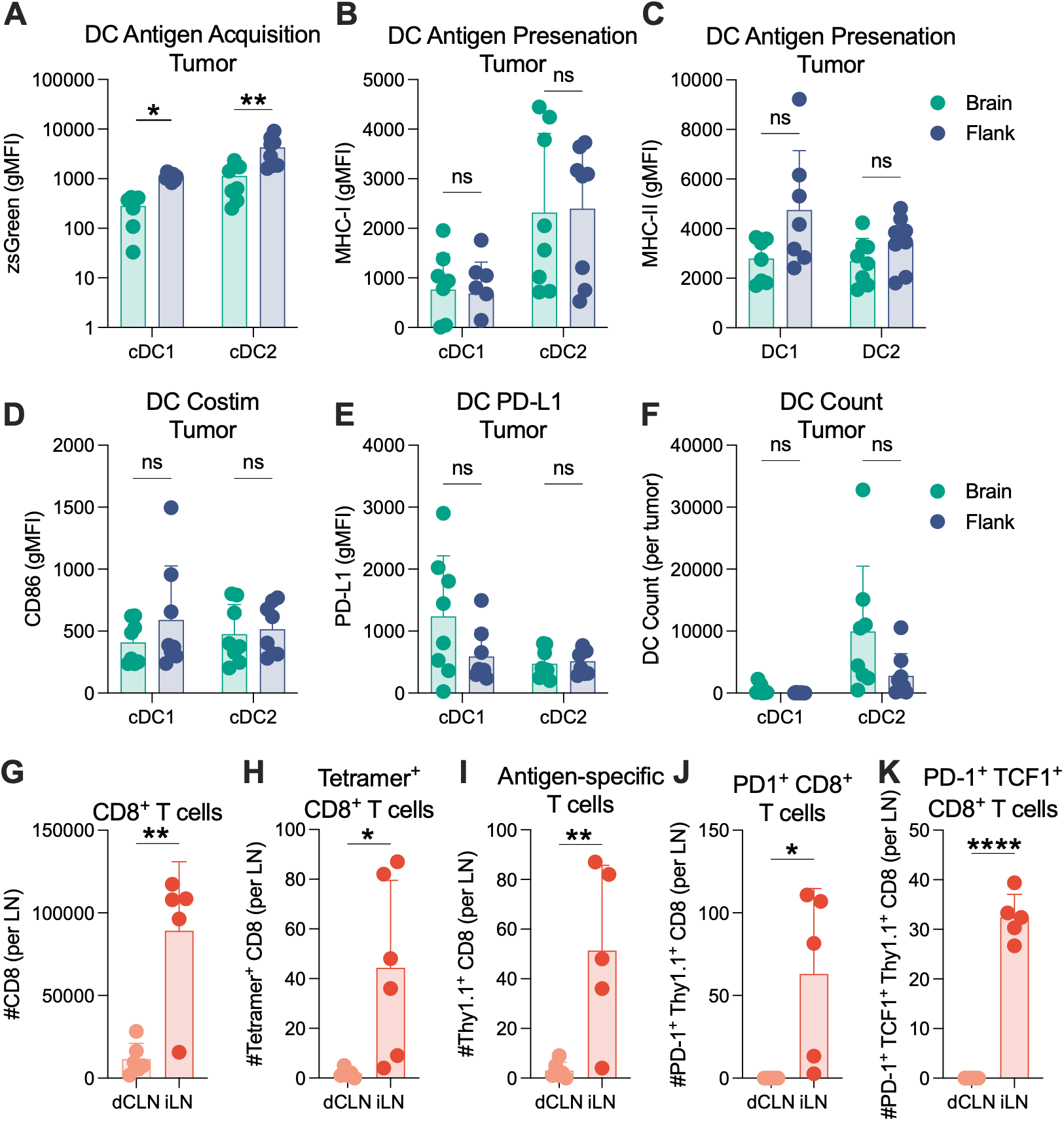
Brain tumors exhibit impaired antigen-specific T cell priming in the tdLN. (A) Quantification of tumor-infiltrating zsGreen^+^ cDC1 and cDC2 in brain or flank tumors 7 days after tumor implantation measured by geometric mean fluorescence intensity (gMFI) (n = 7-8 mice per group; two independent experiments shown). (B) Quantification of tumor-infiltrating cDC1 and cDC2 MHC-I expression in brain or flank tumors 7 days after tumor implantation measured by geometric mean fluorescence intensity (gMFI) (n = 7-8 mice per group; two independent experiments shown). (C) Quantification of tumor-infiltrating cDC1 and cDC2 MHC-II expression in brain or flank tumors 7 days after tumor implantation measured by gMFI (n = 7-8 mice per group; two independent experiments shown). (D) Quantification of tumor-infiltrating cDC1 and cDC2 CD86 expression in brain or flank tumors 7 days after tumor implantation measured by gMFI (n = 7-8 mice per group; two independent experiments shown). (E) Quantification of tumor-infiltrating cDC1 and cDC2 PD-L1 expression in brain or flank tumors 7 days after tumor implantation measured by gMFI (n = 7-8 mice per group; two independent experiments shown). (F) Quantification of tumor-infiltrating cDC1 and cDC2 in brain or flank tumors 7 days after tumor implantation (n = 8 mice per group; two independent experiments shown). (G) Quantification of antigen-specific CD8^+^ T cells in the tdLN. One day before tumor implantation, 5×10^5^ OT-I CD8^+^ T cells were adoptively transferred, and CD8^+^ T cells were quantified in the tdLN 7 days later (n = 5-6 mice per group; one independent experiment shown). (H) Quantification of tetramer-positive CD8^+^ OT-I T cells in the tdLN 7 days after tumor implantation (n = 5-6 mice per group; one independent experiment shown). (I) Quantification of antigen-specific CD8^+^ OT-I T cells in the tdLN 7 days after tumor implantation (n = 5-6 mice per group; one independent experiment shown). (J) Quantification of PD-1^+^ CD8^+^ OT-I T cells in the tdLN 7 days after tumor implantation (n = 5 mice per group; one independent experiment shown). (K) Quantification of PD-1^+^ TCF-1^+^ CD8^+^ OT-I T cells in the tdLN 7 days after tumor implantation (n = 5-6 mice per group; one independent experiment shown). Data are shown as mean ± SD. *p<0.05, **p<0.01, ****p<0.0001, ns: not significant; Student’s t-test (A-K).

**Figure S4.**
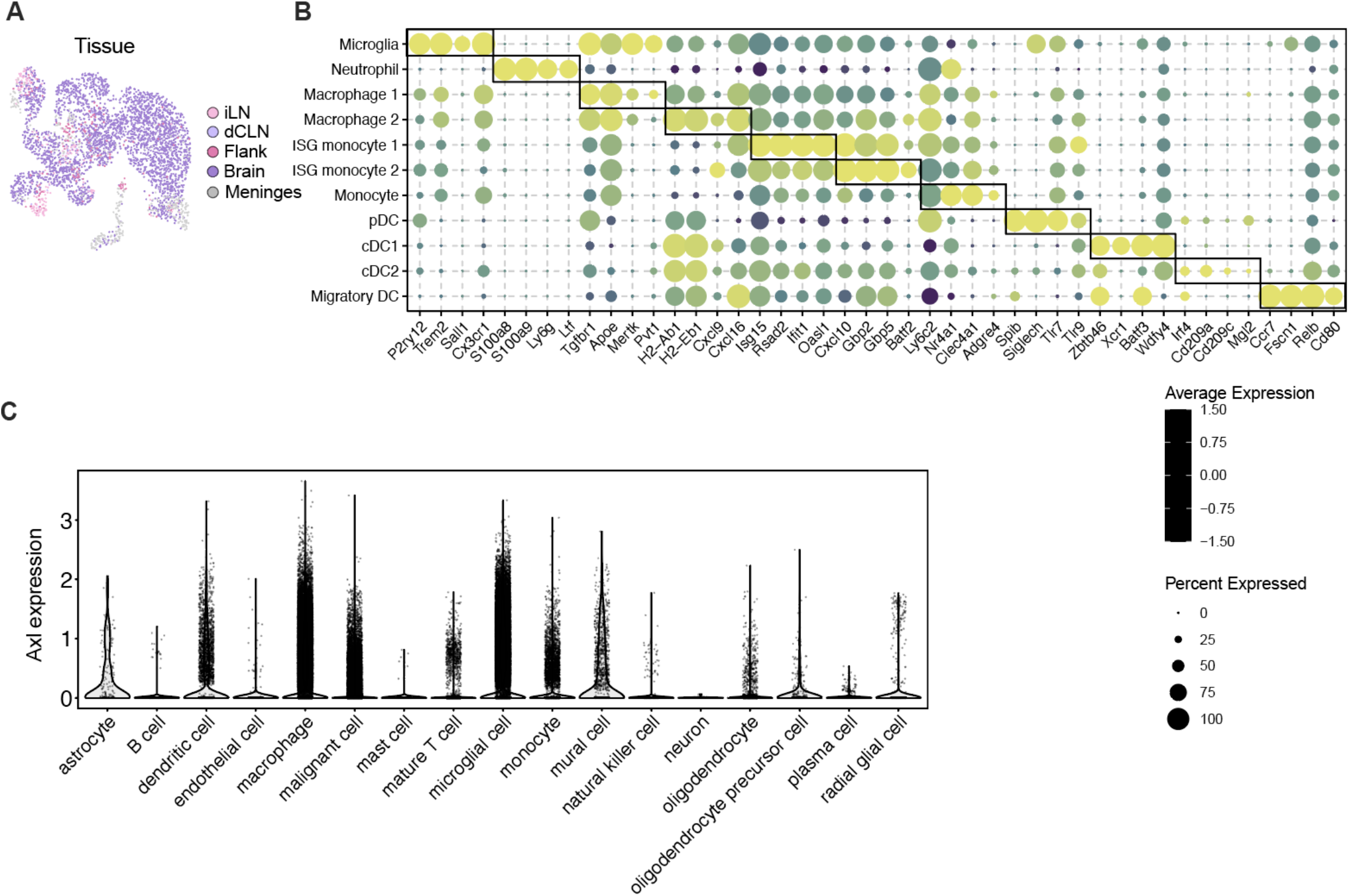
Myeloid cells in the GBM microenvironment. (A) UMAP plot of all myeloid cells analyzed by scRNA-seq, colored by tissue of origin including inguinal lymph node (iLN), deep cervical lymph node (dCLN), flank tumor (Flank), brain tumor (Brain), and meninges (n=3800). (B) Expression of subtype-defining markers. Dot size: percent expressing; color: average expression (n=24,896 cells). (C) Expression of *Axl* in human GBM samples within each of the specified cell types.

**Figure S5.**
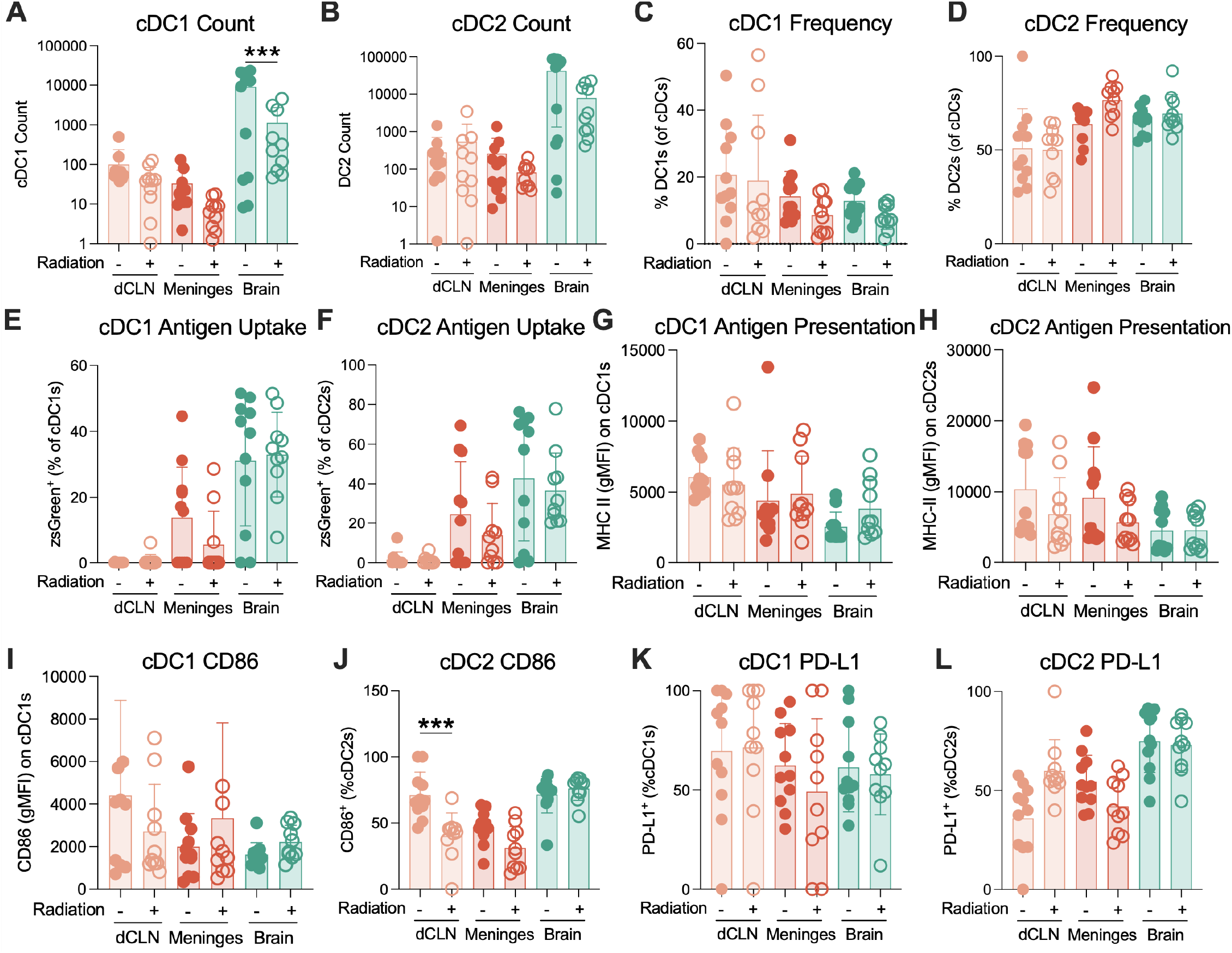
Radiation therapy does not alter classic cDC phenotypic markers. (A) Quantification of cDC1 in deep cervical lymph nodes (dCLN), meninges, or brain tissue from GBM-bearing mice treated with 2×2 Gy RT or sham 14 days after CzO tumor injection. Tissues were harvested 72 hours after completion of RT (n = 10-11 mice per group; two independent experiments shown). (B) Quantification of cDC2 in indicated tissue from GBM-bearing mice treated with 2×2 Gy RT or sham 14 days after CzO tumor injection. Tissues were harvested 72 hours after completion of RT (n = 10-11 mice per group; two independent experiments shown). (C) Quantification of cDC1 frequency in indicated tissue from GBM-bearing mice treated with 2×2 Gy RT or sham 14 days after CzO tumor injection. Tissues were harvested 72 hours after completion of RT (n = 10-11 mice per group; two independent experiments shown). (D) Quantification of cDC2 frequency in indicated tissue from GBM-bearing mice treated with 2×2 Gy RT or sham 14 days after CzO tumor injection. Tissues were harvested 72 hours after completion of RT (n = 10-11 mice per group; two independent experiments shown). (E) Quantification of zsGreen^+^ cDC1 frequency in indicated tissue from GBM-bearing mice treated with 2×2 Gy RT or sham 14 days after CzO tumor injection. Tissues were harvested 72 hours after completion of RT (n = 10-11 mice per group; two independent experiments shown). (F) Quantification of zsGreen^+^ cDC2 frequency in indicated tissue from GBM-bearing mice treated with 2×2 Gy RT or sham 14 days after CzO tumor injection. Tissues were harvested 72 hours after completion of RT (n = 10-11 mice per group; two independent experiments shown). (G) Quantification of cDC1 MHC-II expression by geometric mean fluorescence intensity (gMFI) in indicated tissue from GBM-bearing mice treated with 2×2 Gy RT or sham 14 days after CzO tumor injection. Tissues were harvested 72 hours after completion of RT (n = 10-11 mice per group; two independent experiments shown). (H) Quantification of cDC2 MHC-II expression by gMFI in indicated tissue from GBM-bearing mice treated with 2×2 Gy RT or sham 14 days after CzO tumor injection. Tissues were harvested 72 hours after completion of RT (n = 10-11 mice per group; two independent experiments shown). (I) Quantification of cDC1 CD86 expression by gMFI in indicated tissue from GBM-bearing mice treated with 2×2 Gy RT or sham 14 days after CzO tumor injection. Tissues were harvested 72 hours after completion of RT (n = 10-11 mice per group; two independent experiments shown). (J) Quantification of cDC2 CD86 expression by gMFI in indicated tissue from GBM-bearing mice treated with 2×2 Gy RT or sham 14 days after CzO tumor injection. Tissues were harvested 72 hours after completion of RT (n = 10-11 mice per group; two independent experiments shown). (K) Quantification of cDC1 PD-L1 expression by gMFI in indicated tissue from GBM-bearing mice treated with 2×2 Gy RT or sham 14 days after CzO tumor injection. Tissues were harvested 72 hours after completion of RT (n = 10-11 mice per group; two independent experiments shown). (L) Quantification of cDC2 PD-L1 expression by gMFI in indicated tissue from GBM-bearing mice treated with 2×2 Gy RT or sham 14 days after CzO tumor injection. Tissues were harvested 72 hours after completion of RT (n = 10 mice per group; two independent experiments shown). Data are shown as mean ± SD. ***p<0.001. Two-way ANOVA with Tukey’s multiple comparison test (A-L).

**Figure S6.**
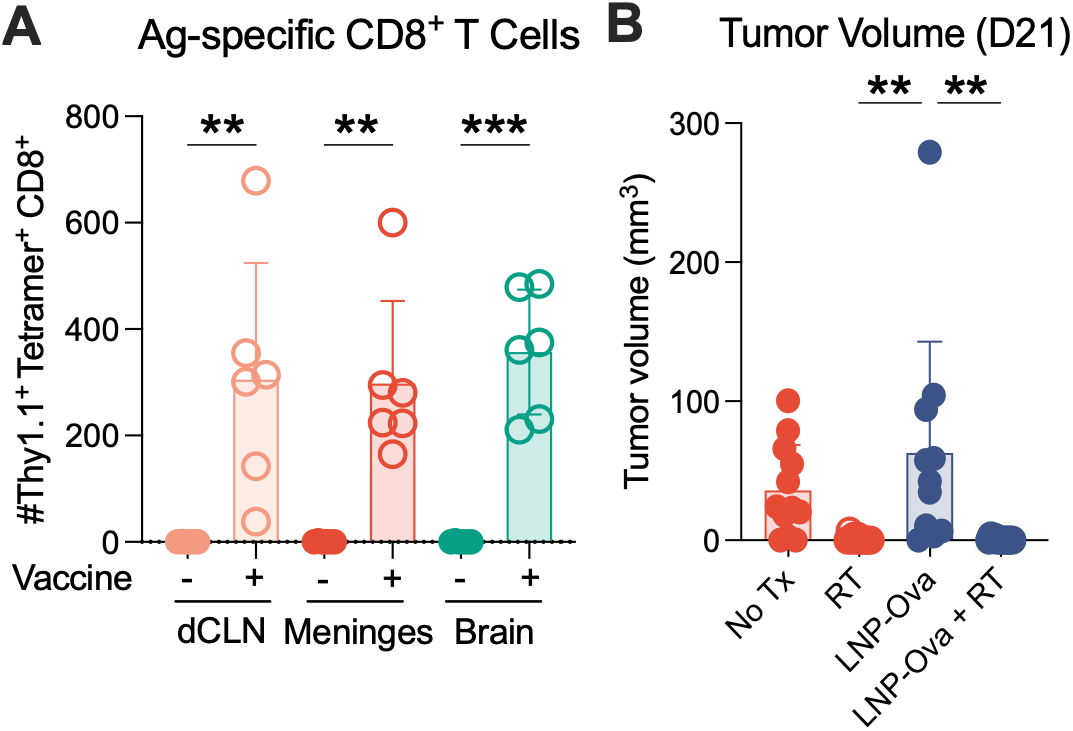
GBM responses to RT and LNP-Ova. (A) Quantification of OT-I^+^ CD8^+^ tetramer-reactive T cells in indicated tissues 21 days after intramuscular LNP-Ova vaccination (n = 5-6 mice per group; one independent experiment shown). (B) Quantification of glioblastoma tumor volume on MRI obtained 21 days after tumor initiation in mice treated with no treatment (No Tx), radiation (RT), mRNA vaccination (LNP-Ova), or mRNA vaccination and radiation (LNP-Ova+RT). Data are shown as mean ± SD. **p<0.01; ***p<0.001. Two-way ANOVA with Tukey’s multiple comparison test (A, B).

**Figure S7.**
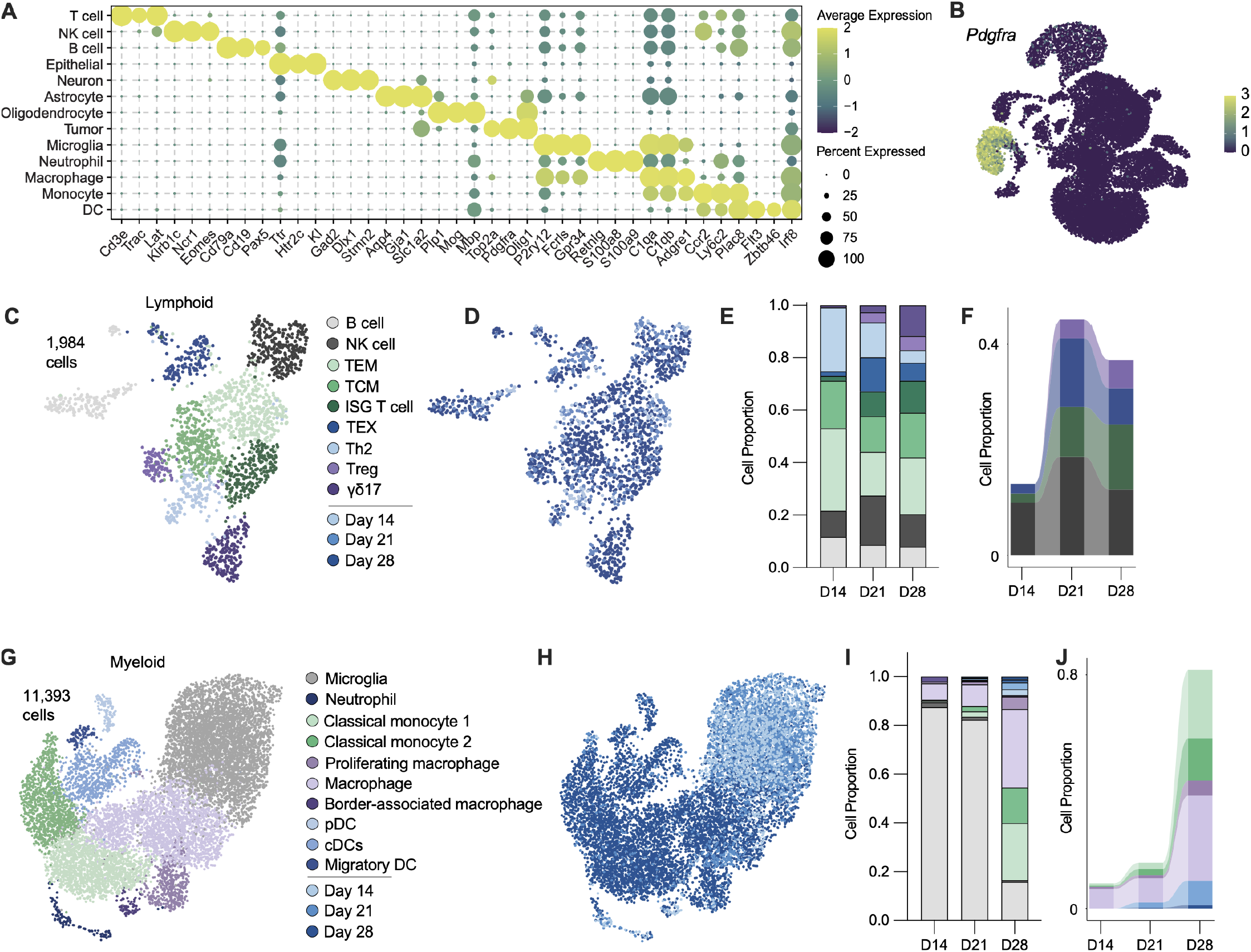
Immune landscape during autochthonous GBM progression. (A) Expression of subtype-defining markers. Dot size: percent expressing; color: average expression (n = 16,076 cells). (B) UMAP plot of all PPzO cells analyzed by scRNA-seq, colored by expression of *Pdgfra* transcript. (C) UMAP plot of all 1,984 lymphoid cells from PPzO tumors analyzed by scRNA-seq, colored by cell annotation including B cells, natural killer (NK) cells, effector memory T cells (TEM), central memory T cells (TCM), T cells with type I interferon-stimulated gene program (ISG T Cell), exhausted T cells (TEX), Th2 T cells, regulatory T cells (Treg), and gamma-delta 17 cells (γδ17). (D) UMAP plot of all 1,984 lymphoid cells from PPzO tumors analyzed by scRNA-seq, colored by date of collection. (E) Proportion of each cell type (annotated in S7C) within each tissue, by time point (F) Proportion of each specified cell type (annotated in S7C) within each tissue, by time point. (G) UMAP plot of all 11,393 myeloid cells from PPzO tumors analyzed by scRNA-seq, colored by cell annotation including microglia, neutrophils, classical monocyte 1, classical monocyte 2, proliferating macrophages, macrophages, border-associated macrophages, plasmacytoid dendritic cells (pDCs), conventional dendritic cells (cDCs), and migratory dendritic cells. (H) UMAP plot of all 11,393 myeloid cells from PPzO tumors analyzed by scRNA-seq, colored by date of collection. (I) Proportion of each cell type (annotated in S7G) within each tissue, by time point. (J) Proportion of each specified cell type (annotated in S7G) within each tissue, by time point.

